# The *Nematostella* synaptonemal complex mediates divergent and sex-specific meiotic programs

**DOI:** 10.64898/2026.01.18.699972

**Authors:** Stefanie Williams, Jennifer Gardner, Stephanie H Nowotarski, Matthew C Gibson

## Abstract

Throughout eukaryotes, the synaptonemal complex (SC) is a supramolecular structure essential for meiotic chromosome dynamics and sexual reproduction. While metazoan SC proteins display significant sequence divergence, the lack of functional analyses beyond bilaterian models has obscured a mechanistic understanding of SC evolution. Here we report that the sea anemone *Nematostella vectensis* exhibits synapsis of homologous chromosomes in a classic zipper-like manner and expresses eight orthologs of ten known vertebrate SC proteins. Surprisingly, mutagenesis of the core SC components *sycp1, syce2*, and *sycp3* resulted in divergent and sexually dimorphic phenotypes, indicating the functional diversification of these conserved factors. Combined, these findings challenge the assumption that the conservation of protein sequence or ultrastructure implies mechanistic homology.

---

Meiosis is a specialized cell division that results in haploid gametes. Early in this process, maternal and paternal copies of each chromosome must pair and genetically recombine to segregate properly (*1*). The synaptonemal complex (SC), a proteinaceous structure that assembles between homologous chromosomes, ensures the formation of crossovers and thus the fidelity of meiotic chromosome segregation (*2*). Indeed, infertility, aneuploidy, and spontaneous abortions in humans have been linked to erroneous SC formation (*3*).

The SC exhibits ultrastructural conservation despite the rapid evolution of its protein components (*4–7*). Consistent with this, *Mus musculus, Drosophila melanogaster*, and *Caenorhabditis elegans* possess SC proteins with highly divergent protein sequences (*4*). Nevertheless, these proteins remain functionally analogous: gross SC protein aberrations lead to failure of chromosome synapsis, crossover defects, and subsequent meiotic arrest and/or chromosome missegregation (*3, 8–17*). Despite the sequence divergence between SC proteins in traditional vertebrate and invertebrate model organisms, Fraune et al. reported that the cnidarian *Hydra vulgaris* expresses five identifiable orthologs of vertebrate SC proteins (*18–20*). While the authors confirmed the morphological conservation of the *Hydra* SC (*18–21*), the genetic requirements for cnidarian SC proteins remain unknown.

As an early-branching phylum phylogenetically positioned between bilaterian animals and non-metazoan organisms, cnidarians hold a crucial position for reconstructing evolutionary events (Fig. 1A). The utility of cnidarian species for evolutionary inference has thus driven the development of extensive genetic and molecular biological tools, positioning several as model organisms. One such model is the sea anemone *Nematostella vectensis* (Fig. 1B) (*22*). *Nematostella* adults are dioecious and can reproduce sexually and asexually through fission (Fig. 1C) (*23–25*). Both sexes have eight internal structures called mesenteries, each of which harbors one gonad (Fig. 1, D-F and fig. S1). Multipotent stem cells drive the continuous production of gametes in adult *Nematostella* (*26*), allowing females to release up to several hundred eggs with a single spawn (*24, 27*). These features make *Nematostella* an ideal model for the functional investigation of the cnidarian meiotic machinery. Here, we analyze the structure of the SC and functionally characterize three of its protein components.

**Fig. 1.**
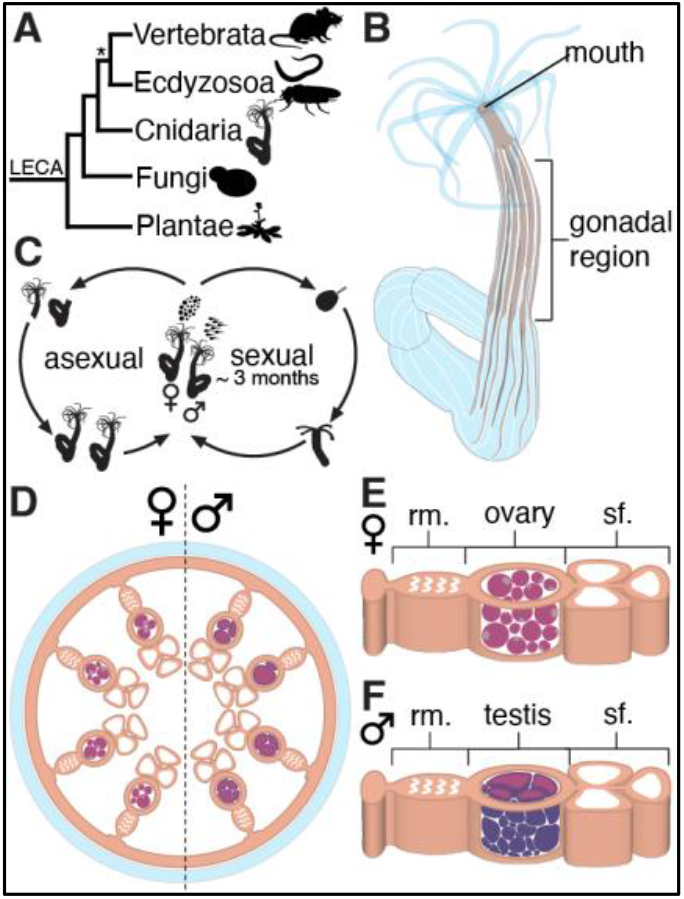
The model sea anemone *Nematostella vectensis*. **(A)** Cartoon of the phylogenetic relationship between *Nematostella* and standard model organisms. *Nematostella* is a non-bilaterian animal, positioned between non-metazoan eukaryotes (such as fungi and plants) and bilaterian animals (such as mice, nematodes, and fruit flies). * denotes the bilaterian lineage. Schematic illustrations of research organisms are from PhyloPic, except for the *Nematostella* illustration, which is from the authors. **(B)** General schematic of the *Nematostella* body. The gonadal region takes up a large portion of the body, and gametes are released through the mouth. **(C)** *Nematostella* adults can reproduce asexually and sexually. **(D)** Schematic cross-section through the gonadal region of a female (left) and male (right) adult. Each adult has eight mesenteries, each containing one gonad. **(E)** Overview of a female mesentery, which generally consists of the reticulate muscle (rm.), the ovary, and the septal filament (sf.). **(F)** Overview of a male mesentery, which, similar to the female one, consists of the reticulate muscle (rm.), the testis, and the septal filament (sf.).

## Results

### Characterizing the *Nematostella* SC

A broadly conserved hallmark of the SC is its ladder-like ultrastructure. To confirm the presence of a canonical SC in *Nematostella*, we analyzed cross-sections of male gonads by electron microscopy. Indeed, by focusing on primary spermatocytes, we observed a typical SC structure (Fig. 2, A to B and table S1). Two electron-dense lateral elements were spaced 98 nm apart (± 13 nm SD) with an electron-dense central element located in the middle (Fig. 2, A to B and table S1). The transverse filaments were less electron-dense (Fig. 2A) and spanned the entire width of the SC (Fig. 2B). With this ultrastructural conservation confirmed, we turned our attention to identifying SC protein components.

**Fig. 2.**
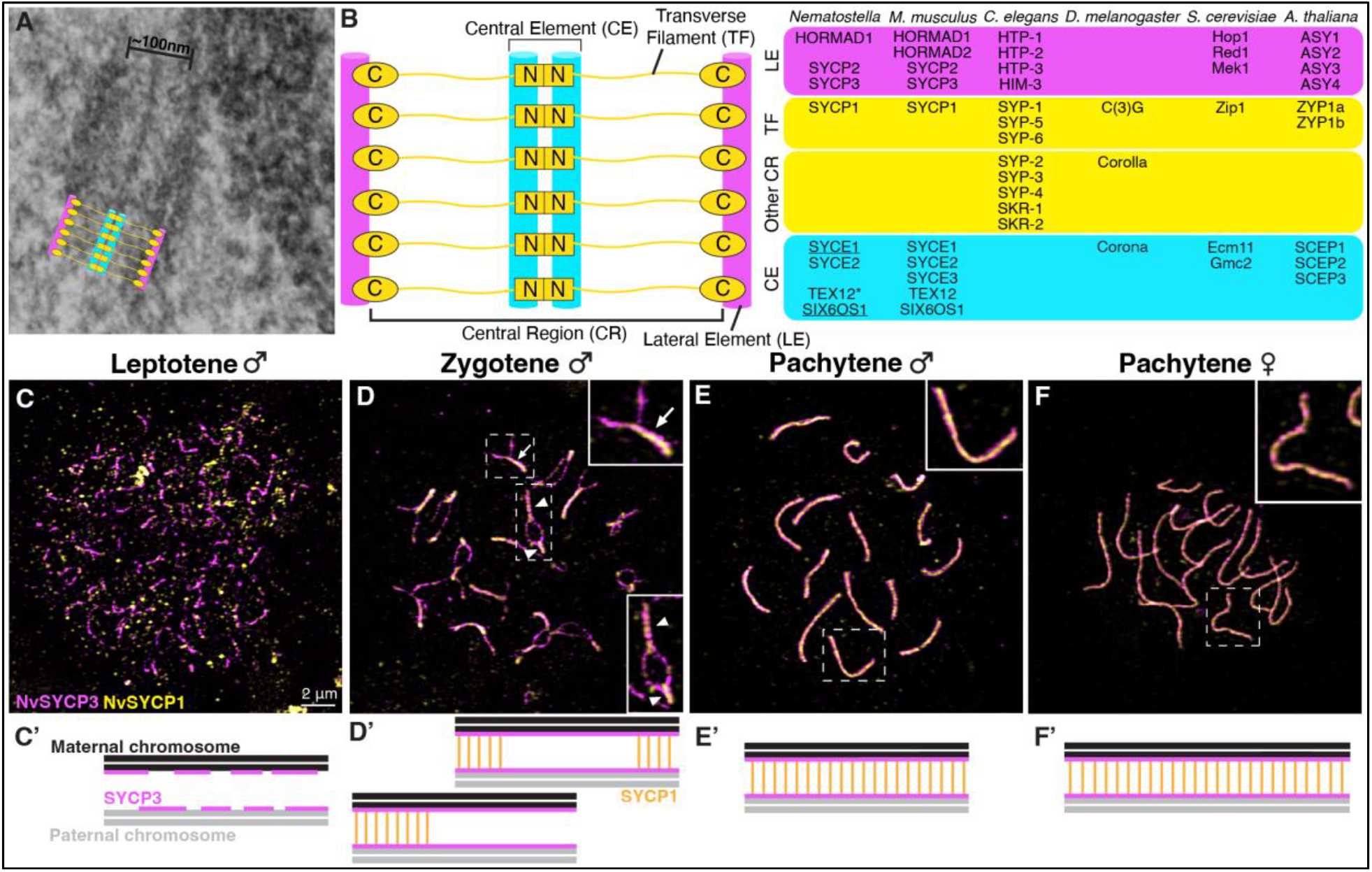
The *wild-type* synaptonemal complex in *Nematostella*. **(A)** Electron microscopy image of a synaptonemal complex (SC) in a *Nematostella* male. The width of the SC is roughly 100 nm. **(B)** The SC consists of two lateral elements (magenta), a central element (cyan), and the transverse filaments that connect these parts (yellow). The protein components that make up these distinct units of the SC are listed from left to right for: *Nematostella vectensis, Mus musculus, Caenorhabditis elegans, Drosophila melanogaster, Saccharomyces cerevisiae*, and *Arabidopsis thaliana*. The underlined *Nematostella* proteins (SYCE1, SIX6OS1) were previously unidentified components of cnidarian SCs. The protein marked with an asterisk (TEX12) had a previously identified *Hydra* ortholog. Note: For simplicity, all cohesin proteins were omitted from the table. **(C)-(F)** Meiotic chromosome spreads from *Nematostella* males (C)- (E) and females (F). Insets highlight example SCs for all relevant stages. All images are at the same scale. Scale bar represents 2 μm. **(D)** The arrow points to synapsis starting from one side while arrowheads point to synapsis initiation from two sides. **(C’)-(F’)** Schematic illustrations of the processes shown in the meiotic chromosome spreads.

Among the five SC proteins previously identified in *Hydra*, four *Nematostella* orthologs were also reported: SYCP1, SYCP2, SYCP3, and SYCE2 (Fig. 2B) (*18–20*). An additional *Nematostella* lateral element component, HORMAD1, was described in 2024 (*28*). Leveraging publicly accessible datasets, we identified *Nematostella* TEX12, previously identified in *Hydra*, and two additional central element proteins: SYCE1 and SIX6OS1 (Fig. 2B, figs. S2-S4). Hence, compared with the vertebrate SC, *Nematostella* lacks only a second HORMAD protein and SYCE3 (Fig. 2B and figs. S5-S11). Notably, the *Nematostella* genome not only encodes a higher number of vertebrate SC orthologs relative to conventional invertebrate models, but the components of each structural element are almost fully conserved. We therefore conclude that the *Nematostella* SC protein repertoire is highly similar to that found in vertebrates. In addition, we identified several orthologs in Porifera and Ctenophora (fig. S5), suggesting their emergence in the metazoan common ancestor.

Immunofluorescence analysis in the freshwater polyp *Hydra* suggests that the cnidarian SC may be functionally equivalent to that observed in bilaterian models (*18–20*). To investigate the localization of *Nematostella* SC proteins, we generated antibodies against the transverse filament protein SYCP1 and the lateral element protein SYCP3. These antibodies were used to stain male and female meiotic chromosome spreads, which were analyzed by Structured Illumination Microscopy. Like in vertebrates, the formation of short SYCP3 threads was first observed in leptotene (Fig. 2, C and C’ and fig. S12). Subsequently, in zygotene, SYCP1 polymerization began at either one or two sites to then continuously synapse the homologous chromosomes (Fig. 2, D and D’ and fig. S12). In pachytene, all 15 chromosome pairs were fully synapsed (Fig. 2, E to F’ and fig. S12). Interestingly, the total pachytene SC length in *Nematostella* females was 1.5 times longer than in males (fig. S12U). Vertebrates like *M. musculus* exhibit similar sex differences: the total SC length in females is twice that of males (*29, 30*). We thus conclude that general SC assembly characteristics and the hitherto unknown mechanisms regulating sexually dimorphic SC lengths are conserved between *Nematostella* and vertebrates.

Vertebrate SC proteins have specific interaction partners that are integral for their structural functions. To further investigate the functional conservation of *Nematostella* SC proteins, we used yeast two-hybrid (Y2H) assays to determine their protein-protein interaction partners. Similar to vertebrate SC proteins, we found that the *Nematostella* proteins SYCP2-SYCP3, SYCE2-TEX12, and SYCE1-SIX6OS1 interacted in 4/4 independent Y2H assays (fig. S13) (*31– 33*). Surprisingly, we observed binding between SYCP1^Cterm^-HORMAD1, suggesting a novel interaction between these *Nematostella* proteins (fig. S13). The C-terminal localization of SYCP1 within the lateral elements makes this possible (*34–36*). Due to the absence of a HORMAD2 ortholog in *Nematostella*, we expected, but failed, to observe an interaction between SYCP2 and HORMAD1 (*31*). Additionally, we were unable to confirm binding between SYCP1 and SYCP2 (*37*). The novel SYCP1^Cterm^-HORMAD1 interaction could explain the absence of these interactions, although negative Y2H results do not necessarily reflect the native context. These assays provide a simplified platform for studying protein-protein interactions that may require post-translational modifications or additional co-factors. Overall, our data suggest broadly conserved interactions between *Nematostella* and vertebrate SC proteins, thereby implying the conservation of their architectural roles. Simultaneously, the SYCP1^Cterm^-HORMAD1 interaction indicates potential functional novelties within the *Nematostella* SC.

### Functional divergence: the sexually dimorphic requirements for *Nematostella* SC proteins

Having established the localization and interaction properties of *Nematostella* SC components, we next systematically investigated the genetic requirement for select SC proteins using CRISPR/Cas9-mediated mutagenesis. We generated small insertion-deletions that introduced premature stop codons in three proteins with architectural roles within distinct structural SC compartments. Specifically, we generated the alleles *sycp1*^*AA1-40*^ to disrupt the transverse filament, *syce2*^*AA1-77*^ to disrupt the central element, and *sycp3*^*AA1-41*^ to disrupt the lateral element (Fig. 2B, Fig. 3, A to C, and fig. S14, A to C). SC proteins contain coiled-coil domains, which are essential for their multimerization and thus their architectural functions (*3*). Accordingly, gRNA target sites were selected to ensure that premature stop codons would preclude translation of coiled-coil domains (Fig. 3, A to C, and fig. S14). In addition, we prevented the translation of the two conserved positively charged domains of SYCP3, which are required for DNA interaction in vertebrates (fig. S14) (*38, 39*). Consistent with a substantial loss of biochemical function, three independent Y2H assays confirmed that the truncated proteins SYCP3^AA1-41^, SYCE2^AA1-77^, and SYCP1^AA1-40^ were unable to bind their interaction partners (fig. S14, E to M).

**Fig. 3.**
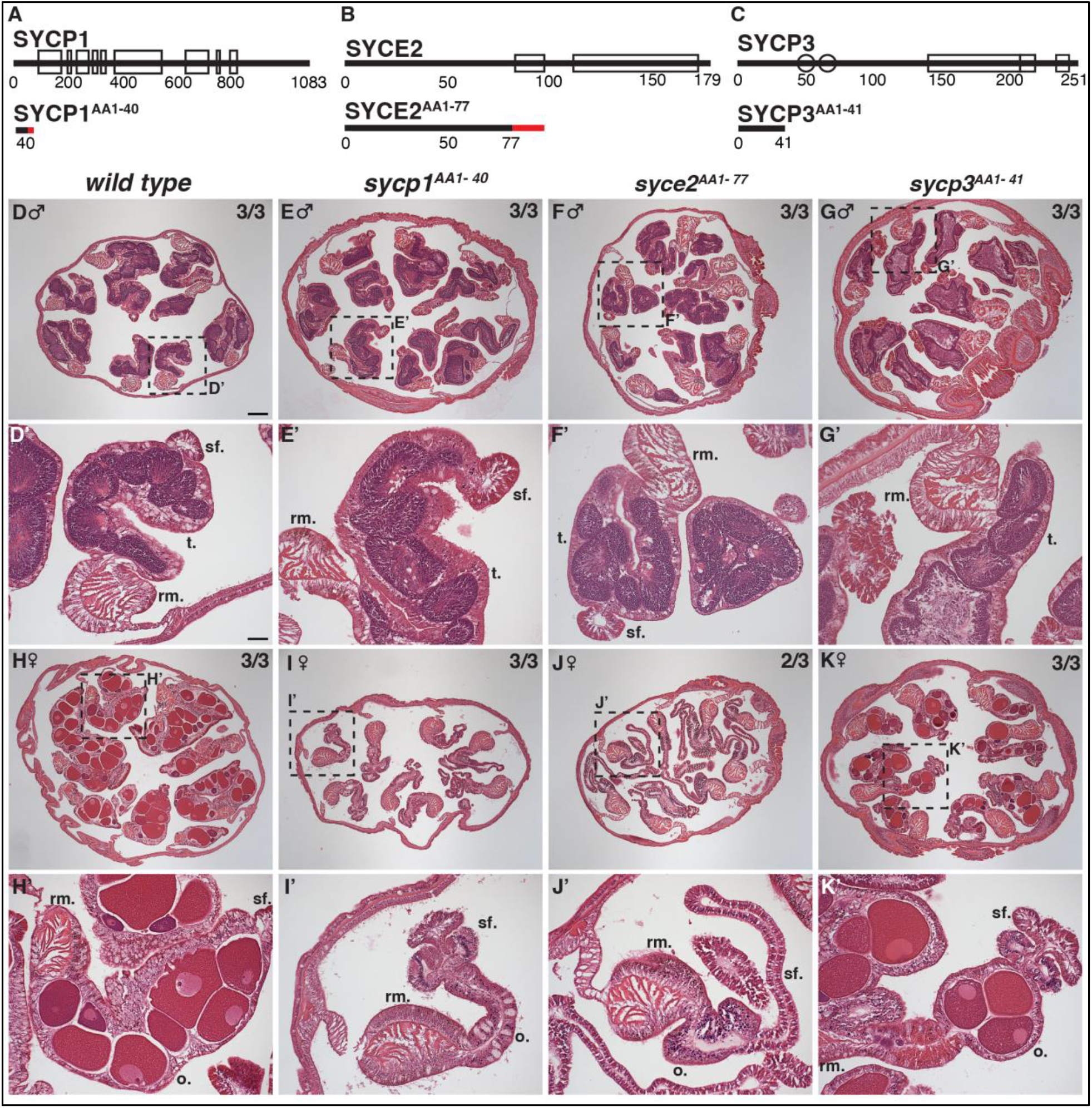
Genetic perturbations of three core SC proteins in *Nematostella*. **(A) - (C)** The top illustration are schematics of the full-length *Nematostella* SC proteins investigated in this study. The black lines represent the entire length of the protein. Numbers below indicate the number of amino acids per protein and their approximate position. Black hollow rectangles signify the locations of coiled-coil domains within each protein. Black hollow circles indicate the approximate locations of positively charged regions found on SYCP3. The illustration below the *wild-type* proteins show the lengths of the truncated proteins in the respective mutants. The red lines indicate amino acids that are translated in the truncated protein but differ from those in the *wild-type* protein due to the insertion-deletion that led to a frameshift and premature stop codon. **(D) - (K)** Images of H&E-stained cross-sections of *wild-type* and mutant gonadal regions of both sexes. Dashed line boxes indicate which region the magnified images in (D’) - (K’) were taken from. Numbers (e.g. 3/3) indicate the number of biological replicates presenting with the shown phenotype out of all biological replicates investigated. Scale bar represents 200 μm. All overview images are at the same scale. **(D’) - (K’)** Magnified images of one mesentery. Scale bar represents 50 μm. All magnified images are at the same scale.

We next established homozygous mutants for each allele and examined the gross morphology of *Nematostella* gonads. Since sexual dimorphisms in meiosis are common (*29, 30, 40*), we separately characterized male and female phenotypes. As expected, we found that *sycp1*^*AA1-40*^ (3/3) and *syce2*^*AA1-77*^ (2/3) homozygous females were sterile. While one of three *syce2*^*AA1-77*^ females had sparsely distributed oocytes within its gonads, there were significantly fewer than observed in controls (Fig. 3, D to K’ and fig. S15). Remarkably, *sycp3*^*AA1-41*^ females and males of all three mutant genotypes were capable of producing gametes (Fig. 3, D toK’ and fig. S15). Spawning data were consistent with this assessment: *sycp3*^*AA1-41*^ homozygous females robustly spawned eggs and homozygous mutant males of all three genotypes successfully released sperm (Table S2). In contrast, even after reaching one year of age, no *sycp1*^*AA1-40*^ females (*n*=6) and just one *syce2*^*AA1-77*^ female (*n*=20) released eggs. In comparison, we observed successful spawning in all *wild-type* and heterozygous sibling controls (Table S2). Thus, we conclude that loss of either SYCP1 or SYCE2 results in female sterility.

To evaluate the structural effects of these mutations on the SC, we prepared chromosome spreads from males homozygous for each of the three alleles. Compared to *wild-type* SCs at pachytene (Fig. 4A and fig. S16), pachytene-like spreads from *sycp1*^*AA1-40*^ males exhibited mostly aligned but asynapsed lateral elements (Fig. 4B and fig. S16). Surprisingly, crossover sites were visible as bridges between the two opposite lateral elements in a total of thirteen spreads from three individual *sycp1*^*AA1-40*^ males (Fig. 4B, fig. S16, and Table S3). While we lack the tools to systematically investigate meiotic crossover formation in *Nematostella*, these data indicate that SYCP1 and synapsis are not essential for crossover formation. In *syce2*^*AA1-77*^ males, all 15 chromosomes were fully synapsed, indicating that SYCE2 is dispensable for *Nematostella* SC assembly (Fig. 4C and fig. S16). Lastly, in pachytene-like spreads from *sycp3*^*AA1-41*^ males, SYCP1 loading appeared unaffected (Fig. 4D and fig. S16). We also detected weak, discontinuous SYCP3 fluorescence signal, suggesting that the truncated SYCP3 N-terminus may interact weakly with the SC (fig. S16). Thus, either a low association of SYCP3 N-termini may allow SYCP1 assembly in *sycp3*^*AA1-41*^ males or the SYCP1^Cterm^-HORMAD1 interaction renders SYCP3 non-essential for synapsis.

**Fig. 4.**
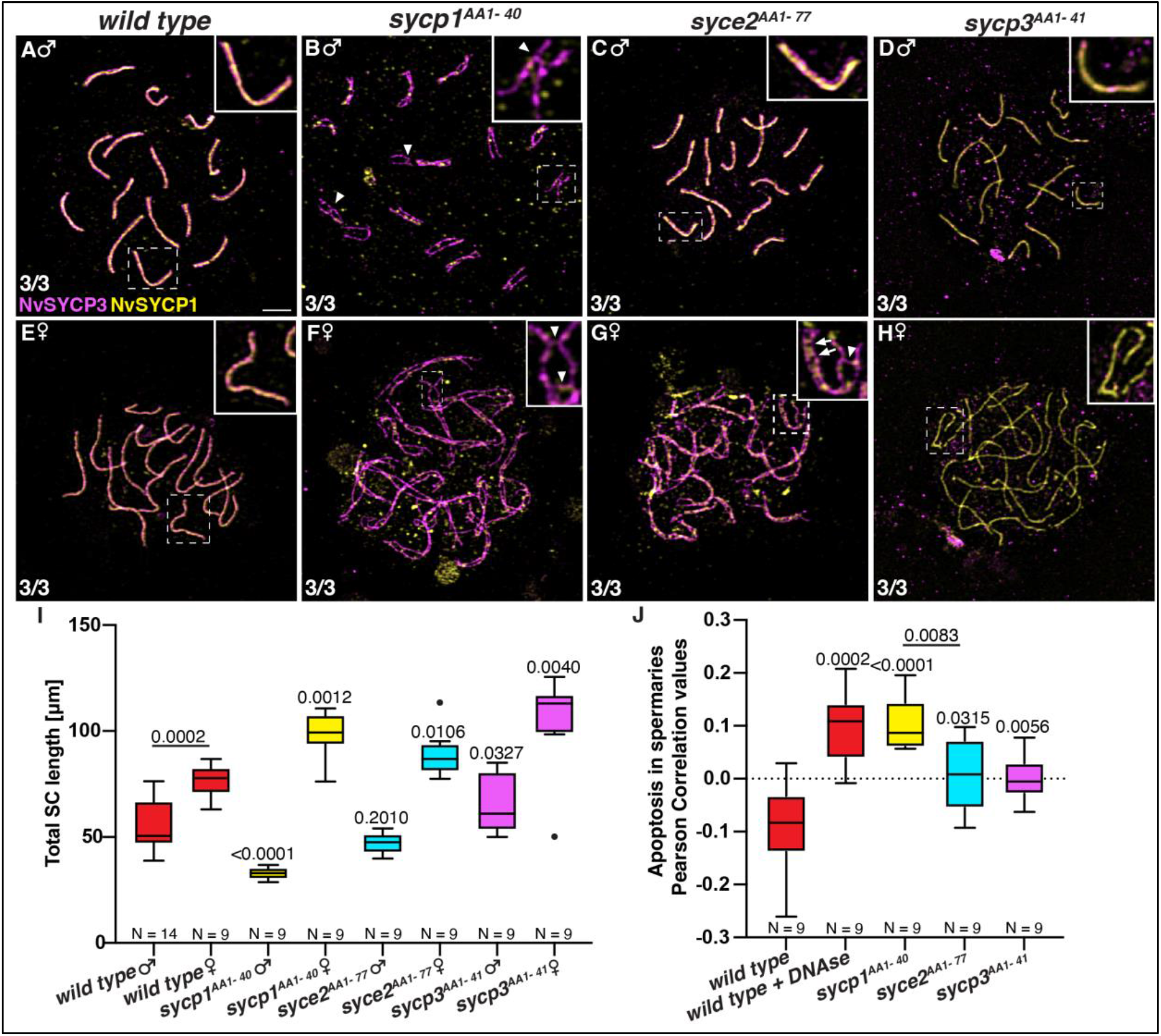
Effects of three genetic perturbations on meiosis in *Nematostella*. **(A) - (H)** Meiotic chromosome spreads from *wild-type* and mutant males and females at pachytene and pachytene-like stages. Insets highlight example SCs for all genotypes; dashed-line boxes indicate their locations in the full image. All images are at the same scale. Scale bar represents 2 μm. Numbers (e.g. 3/3) indicate the number of biological replicates presenting with the shown phenotype out of all biological replicates investigated. **(B)** Arrowheads point to locations of crossovers, visible as bridges formed by SYCP3 of opposite lateral elements. **(G)** Arrows point to locations of gaps in SYCP1 loading and staining between aligned lateral elements. **(I)** Quantification of total SC lengths in all genotypes. P values on top of individual box plots are from a Mann-Whitney test comparing the control total length of the same sex to that of the respective mutant. N indicates the total number of chromosome spreads scored from a total of 3 biological replicates. **(J)** Quantification of TUNEL-positive nuclei relative to all nuclei within the specific ROI in spermaries. P values on top of box plots are the result of a Mann-Whitney test comparing the wild type to the respective treatment or mutant. N indicates the total number of spermaries scored from a total of 3 biological replicates.

We next turned our attention to the female SC. Chromosome spreads from *sycp1*^*AA1-40*^ and *sycp3*^*AA1-41*^ females resembled those of males, showing largely aligned but asynapsed lateral elements in *sycp1*^*AA1-40*^ females, while synapsis was unaffected in *sycp3*^*AA1-41*^ females (Fig. 4, E, F, H and fig. S17). However, the chromosomes in *syce2*^*AA1-77*^ females exhibited discontinuous SYCP1 loading (Fig. 4G, fig. S17), reminiscent of the discontinuous synapsis in male and female *syce2*^*-/-*^ mice (*41*). Similar to asynapsed *sycp1*^*AA1-40*^ males, we observed crossover sites in a total of seven spreads from asynapsed *sycp1*^*AA1-40*^ and four spreads from *syce2*^*AA1-77*^ females (fig. S17, Table S3). Notably, these crossover sites appeared in asynapsed regions of *syce2*^*AA1-77*^ females. These data indicate that SYCE2 is essential for synapsis in females but dispensable in males. Furthermore, the appearance of crossovers in *sycp1*^*AA1-40*^ and *syce2*^*AA1-77*^ females underscores the observation that synapsis is not essential to crossover formation in *Nematostella*.

Upon closer investigation, we found dysregulated SC lengths in most mutants. In particular, *sycp1*^*AA1-40*^ males and females showed opposite phenotypes: SCs were significantly shorter in male mutants but longer in female mutants (Fig. 4I). Given the already sexually dimorphic SC lengths, the *sycp1*^*AA1-40*^ mutation seemingly exacerbates this dimorphism. In addition, SCs were significantly longer in *syce2*^*AA1-77*^ females, as well as *sycp3*^*AA1-41*^ adults of both sexes (Fig. 4I). This suggests a direct involvement of the *Nematostella* SC in regulating meiotic chromosome length. While it is unclear why *syce2*^*AA1-77*^ females exhibit longer SCs, the elongation in *sycp3*^*AA1-41*^ mutants is likely caused by a lack of SYCP3 self-assembly, which requires its coiled-coil domains and has been implicated in chromosome compaction (*38, 39*).

### *Nematostella* SC proteins are not required but advantageous for meiosis

Based on the above observations, we wondered whether mutant meiocytes undergo apoptosis via activation of meiotic checkpoints. To test this, we performed TUNEL assays to detect apoptotic cells in the male gonad (Fig. 4J and fig. S18-19). Low rates of apoptosis were observed in control males, consistent with a functional meiotic checkpoint that prevents defective meiocytes from progressing (Fig. 4J, fig. S18-19). Compared with controls, all homozygous mutant males showed elevated levels of apoptosis (Fig. 4J, fig. S18-19). Moreover, we observed significantly more apoptosis in *sycp1*^*AA1-40*^ males than in *syce2*^*AA1-77*^ or *sycp3*^*AA1-41*^ males. This difference indicates that asynapsis in *sycp1*^*AA1-40*^ males is more disruptive to meiosis than the loss of SYCE2 and SYCP3, which did not perturb male chromosome synapsis. Nonetheless, apoptosis in *syce2*^*AA1-77*^ and *sycp3*^*AA1-41*^ males suggests that SYCE2 and SYCP3 support meiotic processes. As minor changes in SC components can alter SC architecture and dysregulate crossover formation (*42*), *syce2*^*AA1-77*^ and *sycp3*^*AA1-41*^ males may exhibit subtle architectural defects that could lead to aberrant meioses.

To determine the success of meiotic checkpoints, we investigated whether offspring of fertile mutants exhibited increased aneuploidy. We detected no significant aneuploidy in embryos that resulted from the fertilization of *wild-type* eggs with *syce2*^*AA1-77*^ or *sycp3*^*AA1-41*^ males. In contrast, *sycp1*^*AA1-40*^ male offspring showed increased levels of aneuploidy (fig. S20). These findings further support the conclusion that *sycp1*^*AA1-40*^ male meiosis is more dramatically affected than in *syce2*^*AA1-77*^ or *sycp3*^*AA1-41*^ males, coinciding with an inability to eliminate all erroneous meioses. We also investigated the ploidy of embryos from *sycp3*^*AA1-41*^ females, which exhibited increased aneuploidy (fig. S20). Thus, SYCP3 is not essential for meiosis in females but still supports proper chromosome segregation. As aneuploidy differs between *sycp3*^*AA1-41*^ males and females, certain meiotic checkpoints may be differentially sensitive between sexes, as observed in *M. musculus* (*43*). Embryos of the single *syce2*^*AA1-77*^ female that spawned showed no significant aneuploidy, though notably, we were unable to perform experiments in triplicate. Altogether, *Nematostella* can only achieve normal meiotic success with SYCP1, SYCE2, and SYCP3 present.

## Discussion

The ultrastructural conservation of the SC has fascinated scientists since its first description in 1956 (*44, 45*). Considering the rapid evolution of SC protein sequences, their broad functional conservation is similarly compelling (*4–7*). In this first systematic study of cnidarian SC proteins, we demonstrate the coexistence of orthologous relationships with vertebrate proteins alongside dramatically divergent architectural roles. This indicates that conservation of ultrastructure and proteins can coincide with substantial functional divergence.

The dependence of crossover formation and fertility on chromosome synapsis has largely shaped our traditional understanding of SC function (Fig. 5A). While not an intended focus of our study, we unexpectedly observed that *sycp1*^*AA1-40*^ males and *sycp1*^*AA1-40*^ and *syce2*^*AA1-77*^ females were capable of crossover formation between unsynapsed chromosome regions (Fig. 5B). Generally, two distinct crossover pathways exist. These result in class I and class II crossovers, with the latter often considered as back-ups (*46*). Across metazoans, synapsis-deficient individuals exhibit dramatic reductions in crossovers, and most progeny are typically aneuploid (*3, 9, 13–16*). These phenotypes arise from reliance on synapsis-dependent class I crossovers and the inability to rescue them sufficiently via the class II pathway. While *syp-5* mutants in *C. elegans* are capable of some crossover formation, these are dysregulated (*47, 48*). In comparison, synapsis is not required for class I crossover formation in *Saccharomyces cerevisiae* and *Arabidopsis thaliana* (*8, 10–12, 17*). Crucially, the number and distribution of these crossovers are severely dysregulated. Our findings suggest that *Nematostella* is either capable of synapsis-independent class I crossover formation, relies more heavily on the class II crossover pathway, or switches to it more readily than other metazoans. While we currently lack the tools to systematically investigate crossover formation in *Nematostella*, the increased rates of apoptosis in our mutant males, and aneuploidy in *sycp1*^*AA1-41*^ males and *sycp3*^*AA1-41*^ females, suggest an increased occurrence of meiotic errors. Thus, our data strongly indicate dysregulation of crossover formation in all SC mutants, even in those capable of synapsis with incomplete SC protein repertoires.

**Fig. 5.**
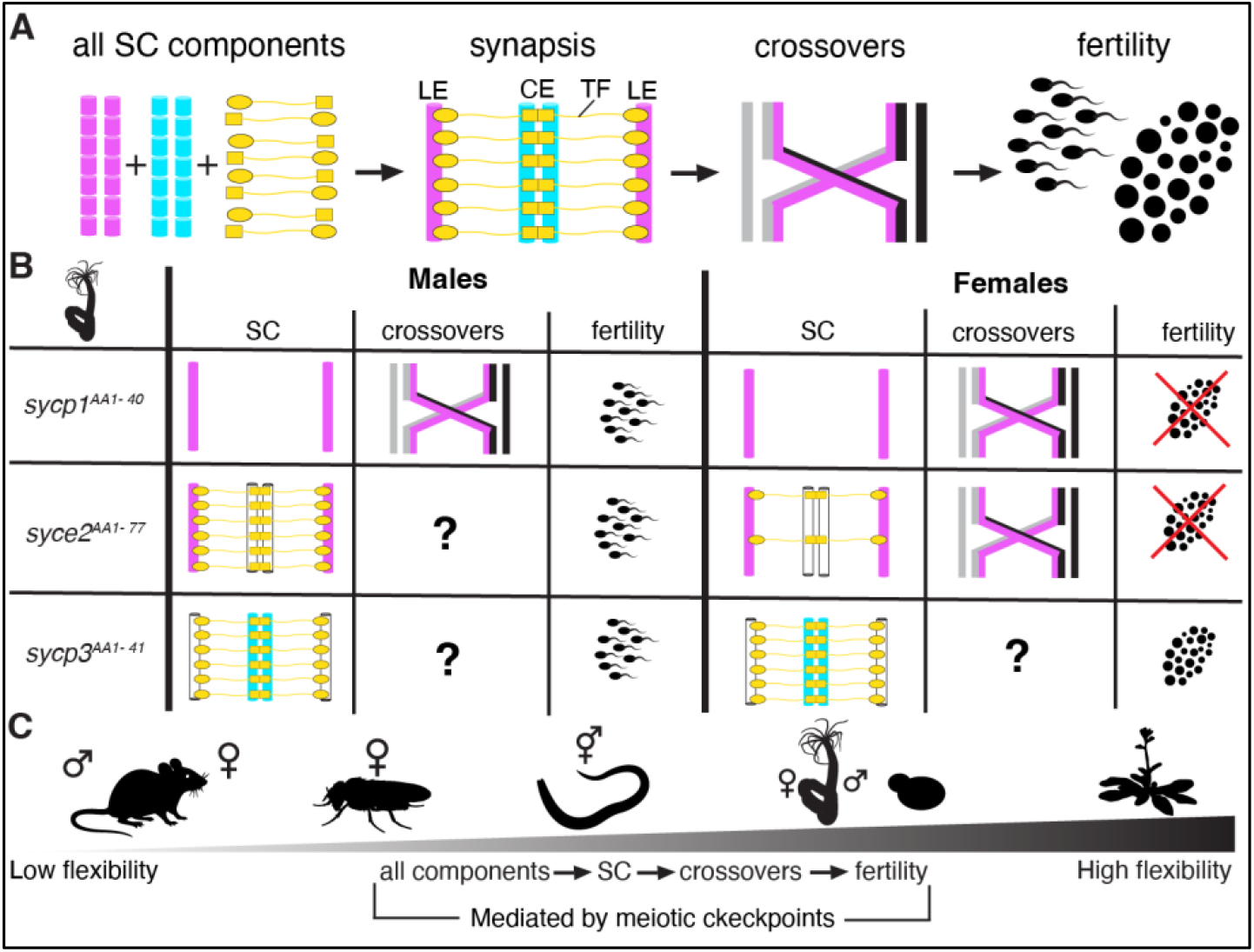
The *Nematostella* SC in relation to other organisms. **(A)** Schematic illustration showcasing that generally all components are required for chromosome synapsis, which in turn is required for crossover formation and, ultimately, fertility. LE = Lateral Element, CE= Central Element, TF = Transverse Filament. **(B)** Schematic illustration of phenotypes observed in the *sycp1*^*AA1-40*^, *syce2*^*AA1-77*^, and *sycp3*^*AA1-41*^ *Nematostella* mutants described here. **(C)** Flexibility of meiotic programs and *Nematostella* phenotypes in relation to SC mutants in standard model organisms used for meiosis research. *M. musculus* mutants exhibit grave SC defects and most often sterility in both sexes. Some female mutants are slightly less severely affected. *D. melanogaster* females exhibit grave SC defects, very low crossover rates and extreme subfertility with offspring highly aneuploid. The same phenotypes apply to *C. elegans* hermaphrodite oogenesis, though some crossovers are achieved in in *syp-5* mutants and, meiosis in *syp-6* mutants is unaffected. Some synapsis deficient *S. cerevisiae* mutants (*ecm11, gmc2*, and *zip1-*N1) exhibit (dysregulated) crossover formation. Many *A. thaliana* SC mutants are synapsis deficient and have (dysregulated) crossovers. The resulting whole picture of the flexibility of meiotic programs is likely mediated by the stringency of meiotic checkpoints.

The observation of synapsis in *sycp3*^*AA1-41*^ mutants, along with the novel SYCP1^Cterm^-HORMAD1 interaction, point to a reshuffling of architectural roles among individual SC proteins and thereby their functional diversification (Fig. 5B). Further architectural differences, such as requiring the structural support provided by SYCE2 (*32, 49*), appear to be sexually dimorphic (Fig. 5B). Given the shorter SCs in *M. musculus* males, Morelli and Cohen previously highlighted the paradoxical nature of their greater structural susceptibility to loss of individual proteins (*30*). We speculate that SC length differences in *Nematostella* may account for the sexually dimorphic requirement for SYCE2 in synapsis. As the execution of downstream processes may be more challenging in longer, unsynapsed SCs, the length differences could also explain the sterility observed in *sycp1*^*AA1-40*^ and *syce2*^*AA1-77*^ females. Alternatively, entry into female meiosis is likely limited due to spatial and energetic constraints of oogenesis. As a result, sterility in *sycp1*^*AA1-40*^ and *syce2*^*AA1-77*^ females may reflect a numerical disadvantage of cells entering meiosis rather than differential effects on meiosis. Overall, meiotic sexual dimorphisms have been extensively documented but remain poorly understood (*29, 30, 40*). Future studies in *Nematostella* may aid investigations as a comparative references.

Despite its functional divergence from vertebrates, the *Nematostella* SC confirms the conservation of core principles: SC proteins assemble into a canonical ladder-like structure and all components are required for the normal completion of meiosis. Previous work confirmed the functional conservation of the DNA double-strand break-inducing enzyme SPO11 in the jellyfish *Clytia hemisphaerica* (*50, 51*). *Clytia spo11* mutants lack SCs and crossovers but remain fertile, producing polyploid embryos (*50*), which may contrast with our findings in *Nematostella*. Apoptosis removes many spermatocytes in *Nematostella* SC mutants. While no data on spermatocyte apoptosis in *Clytia spo11* mutants exist, the robust fertility coinciding with achiasmate chromosomes and polyploid embryos may indicate a weaker meiotic checkpoint. This contrast would be interesting, as the differential sensitivity of meiotic checkpoints most likely determines the flexibility of meiotic programs (Fig. 5C). In comparison to common model species, our data overall highlight that meiotic mechanisms in *Nematostella* are among the most flexible known to date (Fig. 5C). Intriguingly, such differences in meiotic programs may reflect alternative gamete strategies: In *M. musculus*, gamete quality may be strictly controlled because of their limited quantity, especially in females, and the additional energetic investment required by pregnancies. In contrast, a broadcast spawner like *Nematostella* may permit less stringent quality controls due to the high number of developing gametes and absence of post-embryonic maternal investment.

Altogether, our work argues that ultrastructural conservation should not be mistaken for functional invariance. Our findings demonstrate that even core meiotic mechanisms can be evolutionarily reconfigured to achieve alternative functional and sexually dimorphic solutions. By revealing this hidden flexibility within a deeply conserved cellular structure, *Nematostella* emerges as a critical comparative reference point for understanding how evolution preserves SC architecture while rewriting the functions of its constituent proteins.

## Acknowledgments

First, we would like to thank R. Scott Hawley for his invaluable guidance throughout this project until the very day of his passing. As a titan in the pursuit of understanding meiosis and the synaptonemal complex, his support was irreplaceable for the work described. Second, we would like to thank Cathleen M. Lake, Stacie E. Hughes, and Helen R. Horkan for very fruitful discussions and instrumental support throughout the project. Further, we would like to thank Xia Zhao for support with electron microscopy data acquisition, Hannah Wilson for histological sections, and MaryEllen Kirkmeyer, Benjamin Troutwine, Kyle Weaver, and Victoria Hassebroek for support with genetic perturbations. Additional thanks go to Jeff Lange for support with light microscopy. We would like to thank Youbin Xiang and Rachel Nielsen for their support with protein expression and antibody generation. Lastly, we would like to thank Stacie E. Hughes, Helen R. Horkan, Jay R. Unruh, Jennifer L. Gerton, and SaraH Zanders for helpful feedback on the manuscript.

## Author contributions

Conceptualization: SW

Formal analysis: SW

Funding acquisition: MG

Investigation: SW, JG, SN

Supervision: MG

Visualization: SW

Writing – original draft: SW

Writing – review & editing: SW, JG, SN, MG

## Data and materials availability

Original data underlying this manuscript will be made accessible from the Stowers Original Data Repository at https://www.stowers.org/research/publications/libpb-2583

