## supplemental files for "The *Nematostella* synaptonemal complex mediates divergent and sex-specific meiotic programs"

**The PDF file includes:**

Materials and Methods
Supplementary Text
Figs. S1 to S20
Tables S1 to S3

### Materials and Methods

#### Animal care, spawning induction, and sex determination

Generally, *Nematostella vectensis* were kept in 12 ppt artificial sea water (ASW) and fed 2-3 times per week with freshly hatched *Artemia nauplii*. Full water changes were performed the day after feeding. Animals were kept at approximately 27 °C and moved to approximately 18 °C for spawning procedures. Acclimation to 18 °C was allowed for at least one week. Spawning was induced by moving adults kept at 18 °C to room temperature (RT) around 5 PM in front of a turned off light box, covered by a cardboard box to shield the adults from ambient light. At 2 AM, an automated switch turned on the lightbox, providing light exposure until 8:30 AM. Between 8:30 and 9 AM, the water in which the adults were sitting was removed and replaced with water that was kept at 18 °C. Animals were subsequently left at room temperature to spawn, which usually happened around 11 AM. For crossing, egg sac from one individual female was placed into a well containing sperm released into the water by one male. Animals of the F2 generation studied here were fed every two days with full water changes on the days in between.

#### Adult mesentery dissection

Adult animals were left to relax either naturally at RT or by addition of 1:1 MilliQ (MQ) water:12ppt ASW and subsequently paralyzed by addition of 7% Magnesium Chloride Hexahydrate ($MgCl_2$) in 12 ppt ASW. Once paralysis was complete (after approximately 10-15 min), the oral and aboral ends were amputated with a single-use scalpel size 10. With sharp forceps, one mesentery after another was pinched at the aboral opening and pulled out of the adult.

#### Scanning Transmission Electron Microscopy (STEM)

Dissected mesenteries were fixed with 2% paraformaldehyde (PFA), 2.5% gluteraldehyde with 1 mM $CaCl_2$ and 1% Sucrose in 100 mM Na Cacodylate buffer with a pH of 7.38 at room temperature on a nutator for an hour, then moved to 4 °C for overnight. Samples were then rinsed four times for ten minutes each in buffer, then fixed in 1% osmium tetroxide for 2 h at RT. After another set of four buffer rinses, the samples were rinsed in four exchanges of milli-Q water for 10 min each, followed by an overnight incubation in 0.5% aqueous uranyl acetate at 4 °C. The next morning, the samples were rinsed with a set of milli-Q water incubations and subsequently dehydrated in a series of ethanol for 10 min each at the following percentages: 30, 50, 70, 80, 90, 100, 100, 100. Samples were then incubated in Hard Plus Epon resin (Electron Microscopy Sciences) without accelerator at 50% for 5 h, then changed to 100% without accelerator for 48 h at 4 °C, with all steps on a nutator. Three more twelve-hour incubations at 100% Hard Plus resin with accelerator were carried out, and then the embedded samples were polymerized in a 60 °C oven for 48 h. Sections were cut on a Leica UC7 ultramicrotome with a Diatome knife to 80 nm and mounted on a formvar slot grid (Electron Microscopy Sciences). Grids were stained with Sato's lead for 3 min, 1% aqueous uranyl acetate for 4 min, followed by an additional Sato's lead for 5 min. Grids were imaged on a Zeiss Merlin SEM with aSTEM4 detector at 28 kV, 295 pA.

#### X-ray Tomography

The dissected samples from the STEM imaging, in their resin blocks, were used for conventional X-ray tomography imaging. The samples were imaged on a Bruker Skyscan 1272 with a 50 kV X-ray source at 80pA (4W) with no filter, 360 degree rotation at 0.1 degree intervals, 2.25 µm pixel size, with a random movement of 30 and 4 frame averaging. Reconstruction was performed with NRecon software (Bruker, v2.2.0.6), and videos were made with CTvox (Bruker, v3.3.1).

**Development of polyclonal antibodies**

For expression of protein sequences, gene blocks encoding the SYCP1 N-terminus (amino acids 1-210), C-terminus (amino acids 701-910), and full-length SYCP3 plus 6xHis were ordered from IDT. They were cloned into pET-21a(+) vectors and transformed into DH5alpha cells. Single colonies were picked, plasmid purification, and DNA sequence was confirmed using Sanger sequencing. Plasmids were transformed into BL-21 (DE3) cells for standard protein expression followed by protein purification using Ni-NTA His-Bind Resin (EMD Millipore Corp) under denaturing conditions. The purified protein was sent to Cocalico Biologicals, Inc. for injections into rats and guinea pigs. Received total serums were used for antibody purifications: Rat IgG were purified using a HiTrap™ Protein G HP antibody purification column (Cytiva # 29048581) and guinea pig IgG were purified using a HiTrap™ Protein A HP antibody purification column (Cytiva # 29048576) with an ÄKTA pure™ chromatography system. Antibodies were used individually to confirm staining of synaptonemal complexes on surface chromosome spreads.

**Preparation of meiotic chromosome spreads**

The following solutions were prepared in advance: Buffer A (150mM sucrose, 5 mM EDTA, in MQ water, can be stored at 4 ºC), Buffer B (3% PFA, 0.3% Triton-X 100, 150 mM sucrose, 5mM EDTA, in MQ water, must be made fresh on the day of), 0.2% Photoflo (Kodak Photo-Flo 200 solution, Electron Microscopy Sciences #74257). A humid chamber was prepared by placing a wet piece of paper towel into a Corning® 150 mm TC-treated Culture Dish (Corning #430599). Flat tooth picks were placed apart approximately 75 mm on the paper towel. This humid chamber could hold 4 glass slides. Adult mesenteries were dissected out of the animal as described above. Then, mesenteries were placed in buffer A 5 times for 5 min each. For preparation of chromosome spreads, 30 µl of buffer B was placed in the middle of a poly-L-lysine coated microscopy slide (Electron Microscopy Sciences #63410-01). With forceps, one mesentery was placed from buffer A onto buffer B on the slide. With gentle pipetting (set to 10 µl) the mesentery was dissociated. Any larger tissue pieces remaining after gonadal dissociation was complete, were removed from the slide. This was repeated for all mesenteries. Generally, one male mesentery was used on one slide while for females two-four mesenteries were used per slide. The slides were placed onto two flat toothpicks in a humid chamber for 2-3 h. Subsequently, the slides were removed from the humid chamber and left to air dry. Once completely dry, the slides were washed 4 times for 15 min each in 0.2% Photoflo. Slides were left to air dry and then either immediately used for immunofluorescence staining or stored within a slide box at -20 °C for later use.

**Immunofluorescence staining of meiotic chromosome spreads**

The following solutions were prepared in advance: Citrate Buffer, pH 6.0 (0.1M Sodium Citrate, 0.1M Citric Acid, in MQ water), 1xPBS, 1xPBS-T (0.01% Triton-X 100)

If slide was stored at -20 °C, it was moved out to acclimate to RT. Antigen retrieval was then performed by microwaving it in citrate buffer at 70 °C for 15 min total. Afterwards, the slide was moved to 1x PBS. With a kimwipe, a border around the slide was wiped dry and a hydrophobic barrier was drawn with a hydrophobic pen. The slide was then placed onto a slide tray and washed 3 times with 1x PBS-T. Blocking was performed for 1 h at RT in a humid chamber by applying 200 µl of 10% Normal Goat Serum and 3% BSA in PBS-T (0.01% Tween) onto the slide. The blocking solution was removed, and the slide was subsequently incubated with 200 µl of a primary antibody cocktail at 4 °C over night. The next morning, the slide was washed at RT 5 times for 5 min each with PBS-T (0.01% Tween) and then incubated with 200 µl of a secondary antibody cocktail at room temperature for 2 h. Then, the slide was washed again 5 times for 5 min each with PBS-T once, then PBS twice and twice with MQ water. The slide was left to air dry while covered from the dark and then ProLong Glass mounting media was applied to the slide and a coverslip was applied. The mounting media was left to cure for 3 days before microscopy.

**Microscopy of meiotic chromosome spreads**

Structured Illumination Microscopy (SIM) was performed with a 63x, N.A 1.40 Plan-Apo oil objective on a ZEISS Elyra 7 microscope. Alexa Fluor 647 signal was excited with 5% laser power for 100 ms, Alexa Fluor 555 signal was excited with 3% laser power for 80 ms, Alexa Fluor 488 signal was excited with 3% laser power for 50 ms, and DAPI signal (405 nm) was excited with 15% laser power for 100 ms. Images were taken with a z-stack size of 1.0 nm z-stack size and 15 phases. Images were then processed with the SIM tool in the Zeiss software. The parameter was set to "adjust" and the sharpness was set to 10.5 for the channels 647, 568, and 488 and set to 3 for the 405 channel. The method was set to "best fit".

Experiments were done in triplicate. Pachytene-like chromosome spreads were identified as such based on the overall alignment of 15 lateral element pairs for *sycp1*$^{AA1-40}$ and *sycp2*$^{AA1-77}$ mutants. In some instances, alignment was not perfect but achieved for most chromosomes along most of their length. For *sycp3*$^{AA1-41}$ mutants continuous SYCP1 staining of all 15 chromosomes was required.

**Measurements and statistical analysis of total SC lengths and SYCP3 signal intensity on meiotic chromosome spreads**

Images of pachytene and pachytene-like chromosome spreads were max projected. Using the segmented line tool, the entire length of one SC was marked and added to the ROI manager. In case of clear break points, the marking was ended and a new ROI was started. Once all SCs of one spread nucleus were marked and added to the ROI manager, the length of the ROIs were measured using the "Measure" command of the ROI manager. The resulting table was saved as a cvs and the ROIs were saved as a zip file for future reference. For each spread, the total length of all SCs was calculated by summating the length of all SCs of that respective spread. In general, this was done for three technical replicates from three biological replicates at the least. In the case of wild-type males, we measured 14 spreads, each one treated with the same reagents/humidity chamber as the mutant males. All total SC lengths were imported into Prism10 and plotted together in a box plot. Mann-Whitney tests were performed to compare the total SC lengths of mutants to the *wild-type* total SC lengths of the same sex respectively as well as *wild-type* male compared to *wild-type* female total SC lengths.

The same ROIs were used to measure the SYCP3 intensity in *wild-type* and *sycp3* mutant images, however, the line width was increased to 4. A segmented line in the background area was drawn as well. Mean fluorescence intensity was measured across ROIs and the background value was subtracted from all. The average of all these values was calculated, transferred to Prism 10 and plotted as box plots with whiskers. Mann-Whitney tests were used to determine statistical significance of differences.

**Measurements of average line profiles of SYCP1 and SYCP3**

Four images of *wild-type* males and females were used. Per image, 10 straight lines of the same length were drawn perpendicular to two distinguishable later elements of pachytene chromosome spreads. Average line profiles were measured as described previously (*52*, *53*). Some of the total of 40 line profiles had to be discarded due to poor gaussian fits. In the end, the average profiles for anti-SYCP1 and anti-SYCP3 signal were combined, centered, and the raw values of the graphs together with standard deviations were copied into Prism 10 where graphs were plotted.

**Cloning of SC protein coding genes for Yeast Two-Hybrid Assays**

Adult mesenteries from one male were dissected as described above. Mesenteries were placed into 200 µl of TRIzol reagent (ThermoFisher #15596026) and lysed manually with an electric pestle. 200 µl of TRIzol reagent were added and the lysate was moved into a QIAshredder column (Qiagen # 79656) and centrifuged in a tabletop centrifuge at 12000 rpm) for 2 min. 400 µl of 100% ethanol were added to the flow-through and gently mixed with a pipette. The mixture was transferred to two Direct-zol™ RNA MiniPrep (Zymo Research #R2052) columns and centrifuged at 12000 rpm for 30s. The flow-through was discarded and 400 µl RNA Wash buffer was added followed by centrifugation for 30s at 12000 rpm. For the DNAse treatment 5 µl DNAse I were added to 75 µl DNA digestion buffer and the mixture was carefully added onto the column. The DNAse treatment was incubated for 15 min at RT. Then, 400 µl Pre-wash buffer were added, and the columns were centrifuged for 30 s at 12000rpm. The flow-through was discarded and the previous step was repeated. After the flow-through was discarded again, 700 µl of Wash buffer was added and the columns were centrifuged for 2 min at 12000 rpm. The flow-through was discarded. The RNeasy column was placed into an RNAse-free 1.5 ml tube and 30 µl RNAse-free (DEPC-treated) water was added to the column. They were centrifuged for 30 s at 12000 rpm and the tubes were subsequently put on wet ice.

cDNA synthesis was performed either with the iScript™ cDNA synthesis kit (BioRad #1708890) or the SuperScript® III First-Strand Synthesis System for RT-PCR (ThermoFisher #18080-051). For the iScript™ cDNA synthesis kit, 10 µl of the RNA extraction solution were transferred to an RNAse-free tube. The following reagents were added: 4 µl 5x Reaction Mix, 5 µl RNAse-free water, 1 µl Reverse transcriptase. The mixture was incubated in a thermocycler as follows: 5 min at 25 °C, 30 min at 42 °C, 5 min at 85 °C, hold at 4 °C. For the SuperScript® III First-Strand Synthesis System, the manufacturer's protocol was followed, and gene-specific primers were used (Table S3). The DNA concentration was determined using the Qubit™ ssDNA assay kit (ThermoFisher # Q10212) according to manufacturer's recommendations.

DNA amplification was performed with the Phusion™ Plus Green PCR Master Mix (ThermoFisher #F632S) with a 3-step protocol according to manufacturer's recommendations. The DNA was analyzed on a horizontal 1% agarose gel (agarose) with a 1 kb Plus DNA ladder (New England Biolabs # N3200L). DNA fragments at the appropriate size were cut out from the gel and DNA gel purification was performed with the Zymoclean Gel DNA Recovery Kit (Zymo Research #D4002) according to manufacturer's recommendations. pGAD-C1 and pGBDU-C1 yeast two-hybrid vectors (*54*) were linearized with the SmaI restriction enzyme (New England Biolabs #R0141L) according to manufacturer's recommendations in a 60 min reaction. Linearized DNA was analyzed and gel purified as described above. Quantification of DNA concentrations were performed with the Qubit™ dsDNA BR kit (ThermoFisher #Q32853) according to manufacturer's recommendations. Ligation of SC protein coding genes with pGAD-C1 and pGBDU-C1 vectros was performed using the NEBuilder® HiFi DNA Assembly Master Mix (New England Biolabs #E2621L) according to manufacturer's recommendations in a 60 min reaction. Bacterial transformation was performed with 2 µl of the ligation reaction mix using ig® 5-alpha DH5 Competent Cells (Intact Genomics #13-006) according to manufacturer's recommendations. Single colonies were picked and grown in liquid cultures containing 2xYT or LB media with ampicillin added. Liquid cultures were grown over night at 37 °C at 200 rpm. Plasmid DNA purification was performed using the QIAprep Spin Miniprep Kit (Qiagen #27104) according to manufacturer's recommendations. Plasmid DNA was analyzed for positive integration of SC protein coding genes into yeast two-hybrid vectors using Sanger sequencing (service provided by the Stowers Institute for Medical Research) with a primer that amplifies backwards from the multiple cloning site. For positive clones that were longer than Sanger sequencing could resolve, plasmids were submitted to Plasmidsaurus for full-length sequencing to confirm correctness of the sequence. For all truncated versions of SC protein coding genes including the insertion of a superfolder GFP, cloning and verification was performed as described above using plasmids containing the full-length sequences as a template.

Generally, yeast two-hybrid assays were performed as described as previously (*55*). For technical replicates of the assays, matings were newly performed. Four independent experiments were performed for the full Y2H panels and three for the truncated Y2H panels. For the full Y2H panels, haploid cells containing individual binding domain (BD) or activation domain (AD) plasmids were grown in deep well plates overnight at 30 °C in SD-URA or SD-LEU, respectively. The following day, haploids were pinned from liquid cultures onto YPD agar plates using a RoToR HDA pinning robot and long-pin RePads (Singer Instruments). After mating overnight, cells were pinned to SD-LEU-URA agar to select for diploids and allowed to grow for two days. Diploid cells were then robotically inoculated into SD-LEU-URA liquid media, and grown overnight at 30 °C. Cell density was normalized based on OD600, and 5uL was spotted onto SD-LEU-URA, SD-HIS, and SD-HIS + 25mM 3-AT (3-Amino-1,2,4-triazole, Sigma-Aldrich) agar plates using a Biomek i7 Automated Liquid Handler (Beckman Coulter). Plates were grown at 30 °C, and imaged after three, four, and five days using a plate imager (S&P Robotics).

**CRISPR/Cas9-mutagenesis of SC protein coding genes and generation of homozygous mutants**

Mutagenesis of *sycp1*, *syce2*, and *sycp3* in *Nematostella* as well as DNA extraction for genotyping was performed as previously described (*56*). sgRNA sequences were as follows: for *sycp1* 5'- GATCGAGAGTTGAAATGTTG-3', for *syce2* 5'- ACTGGACAGCAGTTTAGTCT-3', and for *sycp3* 5'- CGACGATTTGATGATAGGCG-3'. For sequencing the following primers were used: *sycp1* F1 5'- cactctttccctacacgacgctcttccgatctCTGCAAGTGAAGCAACATAACC-3', *sycp1* R1 5'- gtgactggagttcagacgtgtgctcttccgatctTTCTGCTTCTTGGTGGAGTTT-3', *syce2* F1 5'- CACTCTTTCCCTACACGACGCTCTTCCGATCTctgtcgaggctgttgactatg-3', *syce2* R1 5'- gtgactggagttcagacgtgtgctcttccgatctgatggcccttcatccctattt-3', *sycp3* F1 5'-

cactctttccctacacgacgctcttccgatctTAGTCGCTATGCCTTTGAACC-3’, *sycp3* R1 5’-gtgactggagttcagacgtgtgctcttccgatctTGCAACCGTACGTGAATGT-3’.

F0 animals were reared to adulthood and genetic mutation of individuals was confirmed either using DNA extracts of aboral cuts and/or DNA extracts from embryos that were the result of the F0 crossed to a wild-type individual. Positive F0 adults were crossed to *wild-type* individuals and F1 animals were reared to adulthood. Heterozygosity for genetic mutations was confirmed using DNA extracts of aboral cuts from individuals. Heterozygous animals with the same mutation were crossed to each other to rear a F2 generation to adulthood. All F2 animals were maintained and reared in the same manner to compare *wild-type* to heterozygous to homozygous siblings. Genotypes were determined using DNA extracts of aboral cuts.

**Preparation of cross-sections from *Nematostella* adults**

A 6% PFA solution in ASW was prepared. Adult animals were left to relax either naturally at RT or by addition of 1:1 12ppt ASW:7%MgCl2 and subsequently paralyzed by addition of 7% $MgCl_2$ in 12 ppt ASW. Once paralysis was complete, the oral and aboral ends were amputated. Amputated ends were opened with forceps if needed and the isolated gonadal region was moved into a 1 ml Eppendorf tube containing the 6% PFA solution using a wide bore single use Pasteur pipette. Animals were left to fix at 4 ºC overnight on a nutator. After fixation was complete, animals were washed three times in 1xPBS. For paraffin sections, animals were dehydrated step-wise with ethanol: 10%, 30%, 50%, 70%. Dehydrated animals were stored in 70% ethanol at 4 ºC until paraffin embedding. For cryosections, 1xPBS was removed and replaced with 15% sucrose. Animals were left to settle to the bottom of the tube. Then, the 15% sucrose was replaced with 30% sucrose and again left to settle. Fixed animals in 30% sucrose were stored at 4 ºC until ready for OCT embedding.

For paraffin sections: For paraffin embedding, samples were processed on a Milestone PATHOS Delta Microwave Tissue Processor. The following incubations were performed: 4 min in 70% ethanol, 5 min at 65 ºC in 100% ethanol, 5 min at 45 ºC in isopropyl alcohol, 5 min at 66 ºC in Paraffin Type 9 (Epredia #8337), 5 min at 70 ºC in Paraffin Type 9, 3 min at 70 °C in Paraffin Type 9, 7.5 min at 65 ºC in Paraffin Type 9. Subsequently, the samples were stored at RT in cassettes until ready for embedding in paraffin using a LEICA EG1150H Embedding Center. The samples were sectioned at 7 µm thickness onto charged glass slides (Fisher Scientific #22-042-924) with an Automated Rotary Microtome (HistoCore AUTOCUT #149AUTO00C1 and #14051956472) and an Illuminated Tissue Bath set to 42 °C (230V) (Boekel Scientific #145951-2). Slides were then dried in a dry oven at 36 °C for at least 4 h and then stored at RT until ready for deparaffinzation and staining.

For cryosections: Animals were carefully removed from Eppendorf tubes into a small tray and as much liquid as possible was removed with a kimwipe. A different tray was filled with OCT and animals were carefully moved into OCT. For optimal OCT media penetration, animals were cut into four pieces. Animal pieces were positioned in OCT as desired and subsequently frozen. Frozen tissue was removed from trays and sectioned at 20 µm section thickness. Once completely sectioned, slides were stored at -20 °C until ready for staining.

**Hematoxylin & Eosin (H&E) staining, imaging, and quantification of phenotypes**

Paraffin embedded sections were deparaffinized and stained with H&E as previously described (*57*). Sections were imaged on an inverted Zeiss Axiovert 200 microscope using 5x N.A. 0.16

air, 10x N.A. 0.30 air, 20x N.A. 0.75 air, 40x N.A. 1.3 oil, and 63x N.A. 1.4 oil objectives after Köhler illumination. µManager software was used to acquire images (*58*, *59*). White balance correction was performed in FIJI 2.16.0/1.54p using a publicly available macro (*60*).

Phenotypes of egg and sperm production in controls and mutants were performed using 3 images from 3 biological replicates. For egg scoring, all eggs found in these images for control and mutant females were scored. Mann-Whitney tests were performed to determine statistical significance of egg number differences.

**Immunofluorescence staining and imaging of cross-sections**

The following solutions were prepared in advance: Citrate Buffer, pH 6.0 (0.1M Sodium Citrate, 0.1M Citric Acid, in MQ water), Quenching solution (0.1M Glycine in 1x PBS), High-salt PBS (500 mM NaCl in 1xPBS), 1xPBS-T (0.5% Triton-X 100). Paraffin sections were deparaffinized using the following protocol: Deparaffinization was followed by antigen retrieval with Citrate Buffer at 70 ºC for 15 min in an EZ retriever v3 microwave. Slides were then moved to 1xPBS. We applied 1xPBS to a sequence coverplate and "mounted" the slides to it. Holding the coverplate and slide together, they were added to a manual sequenza staining rack. The slides were washed three times with 1xPBS. To quench free aldehyde groups, quenching solution was applied and incubated for 10 min at RT. Slides were washed three times with 1xPBS. Slides were then permeabilized with 1xPBS-T for 30 min at RT followed by blocking with 10% normal goat serum and 1% BSA for 45 min. Primary antibody solution was prepared in 1x PBS-T (200 µl per slide) containing 3% normal goat serum and 1% BSA. Primary antibody solution was applied to slides and incubated overnight at 4 ºC. Slides were then washed 5 times for 5 min each at RT and incubated with secondary antibodies in 1xPBS-T for 1 h at RT. Then, slides were washed once with 1x PBS, once with high-salt PBS, once more with 1xPBS, and twice with MQ water. Slides were left to air dry and the mounting media of choice was applied (ProLong Gold for standard confocal microscopy, ProLong Glass for SIM) and left for at least 24 h to cure.

**TUNEL staining and confocal microscopy of cross-sections**

Experiments were done in triplicate. The Click-iT™ Plus TUNEL Assay Kits for In Situ Apoptosis Detection (ThermoFisher #C10618) was used for TUNEL staining. The manufacturer's protocol was used with some changes. Stock solutions were prepared as described in the manufacturer's protocol. For the purpose of TUNEL staining on *Nematostella* cross-sections from gonadal regions, only cryosections can be used. This protocol was optimized for 20 µm thick cryosections. Cryosections were moved from -20 °C to RT and left to acclimate. 1xPBS was applied to a sequenza coverplate and a slide containing cryosections was "mounted" to it. While holding the coverplate and slide together, they were placed into a sequenza manual staining rack. The slides were washed twice in 1xPBS until all of the liquid ran through the capillary chamber. Freshly prepared 4% PFA was applied onto the slides and sections were incubated in it for 15 min at RT. Then, the slides were washed with 1xPBS twice for 5 min. 200 µl of Proteinase K solution was applied to each slide which were left to incubate for 30 min at RT. The slides were washed once with MQ water. For a positive control with guaranteed DNA double-strand breaks, a mixture of 1 unit DNAse I in 100 µl 1x DNAse buffer (ThermoFisher #18068015) per positive control slide was prepared. 100 µl of this mixture were added to the positive control slides (in our case: three biological wild-type replicates) and left to incubate for 30 min at RT. The slides were then washed with MQ water once. 100 µl of TdT Reaction buffer was added to each slide and then incubated for 10 min at 37 °C. In the meantime, 100 µl TdT reaction solution was prepared: 94 µl TdT Reaction buffer, 2 µl EdUTP, 4 µl TdT enzyme. 100 µl of this reaction solution was added to each slide which were then incubated for 60 min at 37 °C. Following this, the slides were washed once with MQ water, then blocked with 3% BSA in 1xPBS-T (0.5% Triton X-100) for 5 min. The slides were washed once with 1xPBS. In the meantime, a 10x Click-iT Plus TUNEL Reaction Buffer Additive was prepared by diluting the 100x stock-solution 1:10 in MQ water (the 10x solution needs to be prepared freshly on the day of). Then, 100 µl per slide of the Click-iT Plus TUNEL reaction cocktail was prepared: 90 µl Click-iT Plus TUNEL supermix, 10 µl 10x Click-iT Plus TUNEL Reaction Buffer Additive, mixed by vortexing. This reaction cocktail has to be used within 15 min. 100 µl of this cocktail was added onto the slides which were incubated for 30 min at 37 °C protected by light. From this point, slides should be protected by light whenever possible. The slides were washed with 3% BSA in 1xPBS for 5 min followed by a wash with 1xPBS. Then, Hoechst 34580 (5mg/ml in MQ) (ThermoFisher #H21486) was diluted at 1:500 in 1xPBS and 200 µl were added to each slide and left to incubate for 30 min at RT. The slides were washed twice with 1xPBS and twice with MQ water. Then, MQ water was added onto the slides to carefully remove the sequenza coverplate. The slides were left to air dry and mounted in ProLong Gold (ThermoFisher #P36930). The mounting media was left to cure for at least 24 h prior to imaging.

TUNEL and DNA staining was imaged on a Zeiss LSM980 with a 5x N.A. 0.25 air, and 63x N.A. 1.4 oil objectives. The power of the 405 nm, 488 nm, 561 nm, and 639 nm lasers were set to 1.0%. For images acquired with the 5x objective, the 405 nm and 561 nm lasers were set to a master gain of 650 V each. For images acquired with the 63x objective, the master gain of the 405 nm laser was set to 450 V, 550 V for the 488 and 561 nm lasers, and 650 V for the 639 nm laser. Top and bottom of the section were selected and optimal z stack step sizes were used for image acquisition.

**Quantification of TUNEL positive nuclei**

Three images from three biological replicates were analyzed. The acquired z-stacks were opened in FIJI and the 405 (UV) and 568 (red) channels of the middle z-section were duplicated. Based on the UV channel alone, a region of interest (ROI) was defined using the polygon selection tool. The ROI was roughly drawn to exclude the 2n 2c (meiosis I) nuclei of spermaries. The ROI was added to the ROI tool before the two channels were split. Then, the ROI was added to the red channel and the FIJI plugin “colocalization test” was opened. The channels were selected accordingly and the ROI was defined as being on the red channel. The plugin was run and the resulting cvs table as well as the ROI were saved for each analyzed image. Once all images were analyzed in the same manner, the pearson correlation coefficients were transferred to Prism 10. Because a gaussian distribution could not be assumed, a Mann-Whitney test was performed between the individual genotypes as well as between the wild-type and the DNAse treated control. Pearson correlation coefficients were visualized using box plots with whiskers with the tukey setting, displaying outliers as individual points.

**Preparation, staining, microscopy, and quantification of mitotic chromosome squashes**

For analysis of numerical chromosome aberrations, the eggs and sperm from mutants were used to fertilize with wild-type gametes in individual wells of a 12-well plate. Embryos were left to develop in egg sacs until reaching approximately the 64- and 128-cell-stage. From here, mitotic chromosome squashes were performed as described previously (*61*) with minor changes: after fixation, egg sacs were gently dissociated by carefully pipetting with a P-200 set to 200 µl. This led to some loss of cell-cell-adhesion of embryos but chromosome squashes could still be performed. Once ready for staining, the slides were washed with three times with 1x PBS. Then, Hoechst 34580 (5mg/ml in MQ) (ThermoFisher #H21486) was diluted at 1:500 in 1xPBS and 200 µl were added to each slide and left to incubate for 30 min at RT. Slides were washed three times with 1xPBS, and twice with MQ water. The slides were left to air dry and mounted in ProLong Gold (ThermoFisher #P36930). The mounting media was left to cure for at least 24 h prior to imaging. DNA staining was imaged on a Zeiss LSM980 with a 63x N.A. 1.4 oil objectives. The power of the 405 nm laser was set to 1.0% and the master gain was set to 450 V. Top and bottom of the section were selected and optimal z stack step sizes (0.16 µm) were used for image acquisition.

Max projections of acquired images were performed and numbers of chromosomes were scored. For each genotype, at least 3 images from 3 biological replicates were scored except for the female *syce2* parent where only one single individual spawned one time. Numbers of scored chromosomes were transferred to Prism 10 and plotted as box plots with whiskers. To compare the variance of the data, which would reveal significant increases in aneuploidy, F tests were performed.

**Figure preparation of microscopy images**

Max projections of images were performed, and a region of interest was duplicated. For merged images, channels of interest were merged. Look up tables for CMYK Magenta and CMYK yellow were applied to SYCP3 and SYCP1 channels. Only minimal changes with the brigtness/contrast tool were performed. Images were copied to Adobe Illustrator and adjusted in size while maintaining the original image ratio. For insets, the image was duplicated in Adobe Illustrator and the region of interest was cropped out.

**Ortholog identification**

For the identification of synaptonemal complex protein orthologs in *Nematostella* and other animal groups, confirmed sequences were used for blastp searches with the algorithm BLOSUM45 in the uniprot database, the NCBI protein database, and tblastn searches in the NCBI nucleotide database. Found sequences were tested for presence of coiled-coil domains with DeepCoils2 (*62–64*) and known synaptonemal complex protein related domains using InterPro scan (*65*). Then, sequences were aligned to known orthologs in Geneious Prime version 2024.0.5 using MUSCLE alignment, the PPP algorithm. Once a confident hit was found, they were used for further search in animal groups more basal to the identified protein.

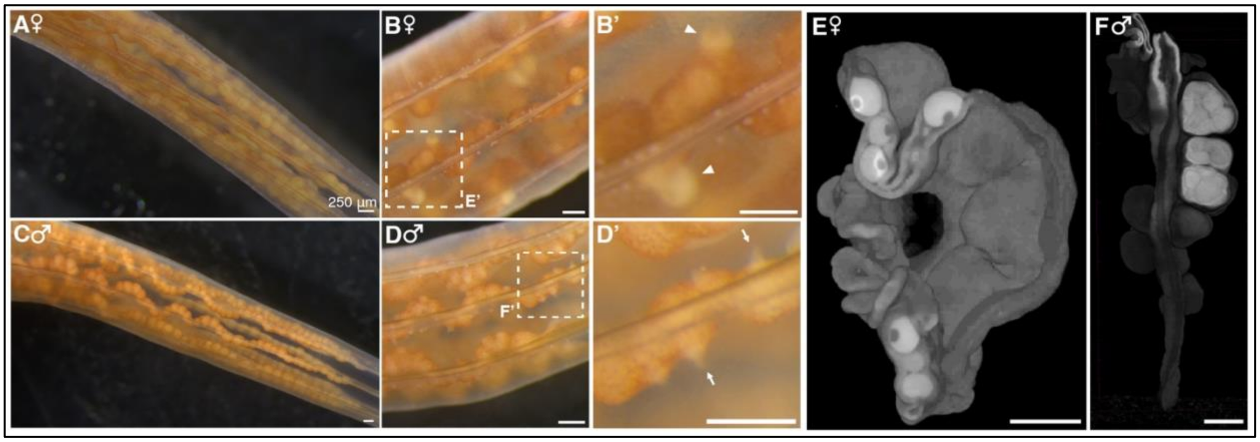


**Fig. S1. The male and female gonads in *Nematostella*.**

**(A)** Photograph of a female gonadal region. The eggs within the gonads are visible. **(B)** Close up of a female gonadal region in which eggs are released from the gonad into the body lumen. Dashed line box indicates where the magnified picture in (B') is from. **(B')** Magnified picture showing eggs in the process of being released from the gonad of a female into the body lumen. Arrowheads point to individual eggs being released. **(C)** Photograph of a male gonadal region. **(D)** Close up of a male gonadal region in which sperm is released from the gonad into the body lumen. Dashed line box indicates where the magnified picture in (D') is from. **(D')** Magnified picture showing sperm release from the gonad of a male into the body lumen. Arrows point to individual locations where sperm is released from. **(E)** X-ray tomography of a female mesentery, digitally sectioned through the gonad to visualize developing oocytes. Oocytes of similar sizes can be found throughout the female gonad without apparent spatiotemporal localization differences. **(F)** X-ray tomography of a male mesentery, digitally sectioned through the gonad to visualize spermaries which are visible as individual "sperm bundles". (A) - (F) Scale bar represents 250 µm.

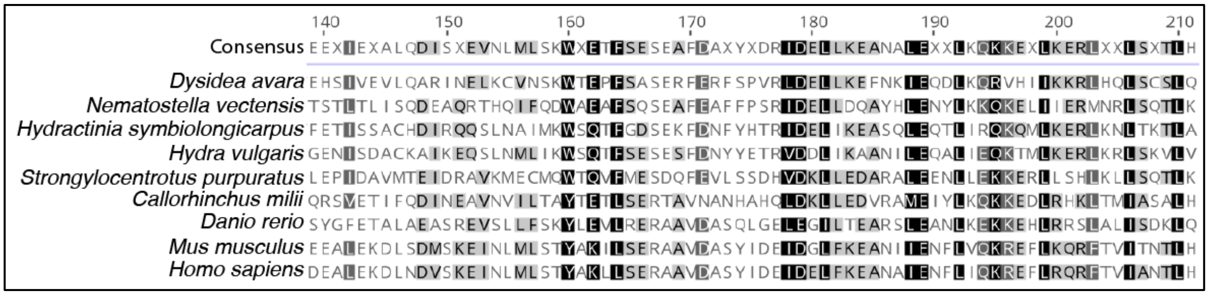


**Fig. S2. Alignment of TEX12 region from various metazoan species.**

Multiple sequence alignment of central region of TEX12 sequences. Black shading of amino acids indicates 100% residue similarity. Dark grey shading of amino acids indicates 80-99% residue similarity. Light grey shading indicates 60-79% residue similarity. White shading indicates less than 60% similarity.

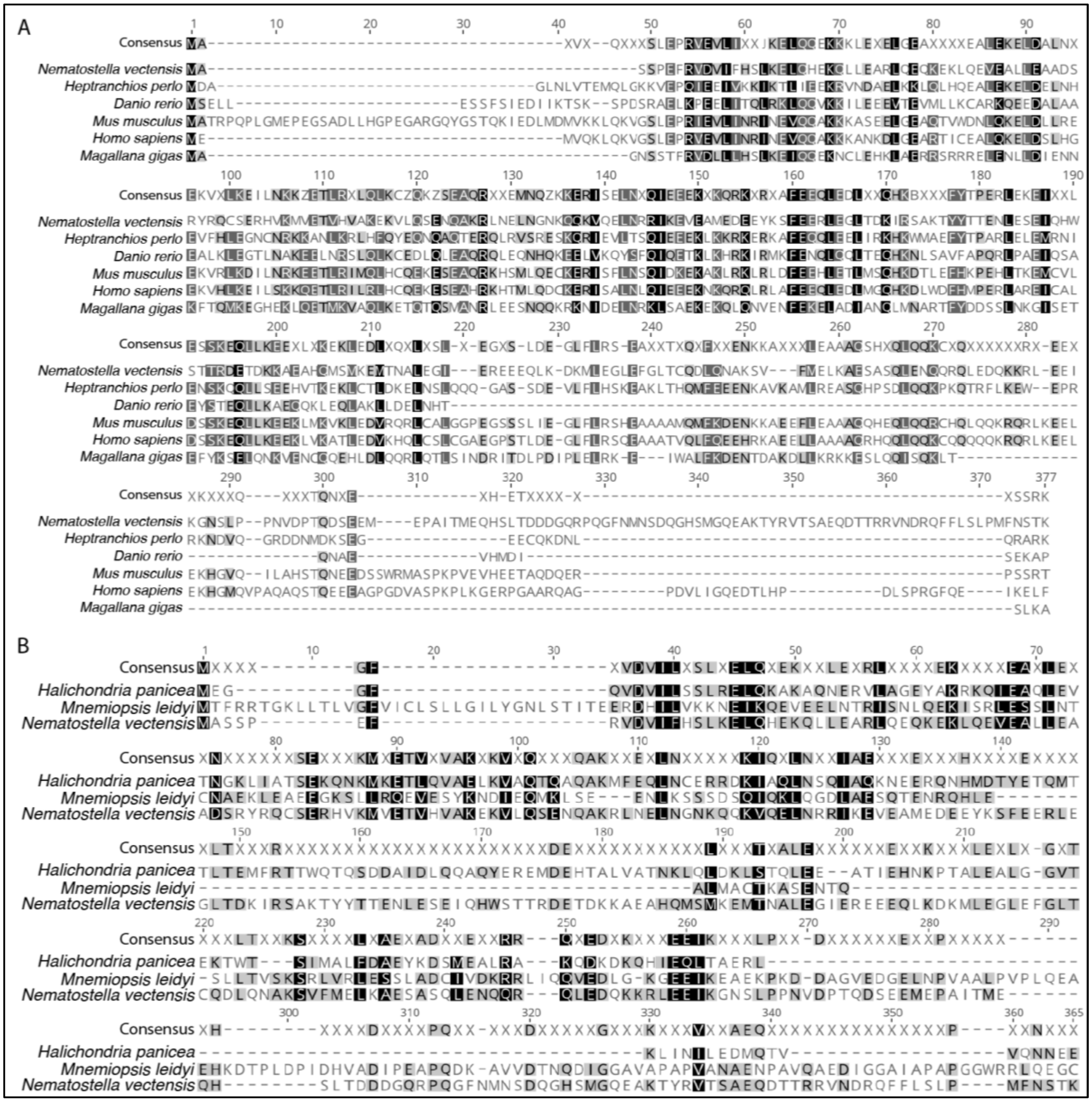


**Fig. S3. Alignment of conserved SYCE1 sequences from various metazoan species.** Multiple sequence alignment of full SYCE1 sequences. Black shading of amino acids indicates 100% residue similarity. Dark grey shading of amino acids indicates 80-99% residue similarity. Light grey shading indicates 60-79% residue similarity. White shading indicates less than 60% similarity. **(A)** Sequences ranging from cnidarians to vertebrates and lophotrochozoans. **(B)** Sequences from a Porifera, a Ctenophora, and *Nematostella*.

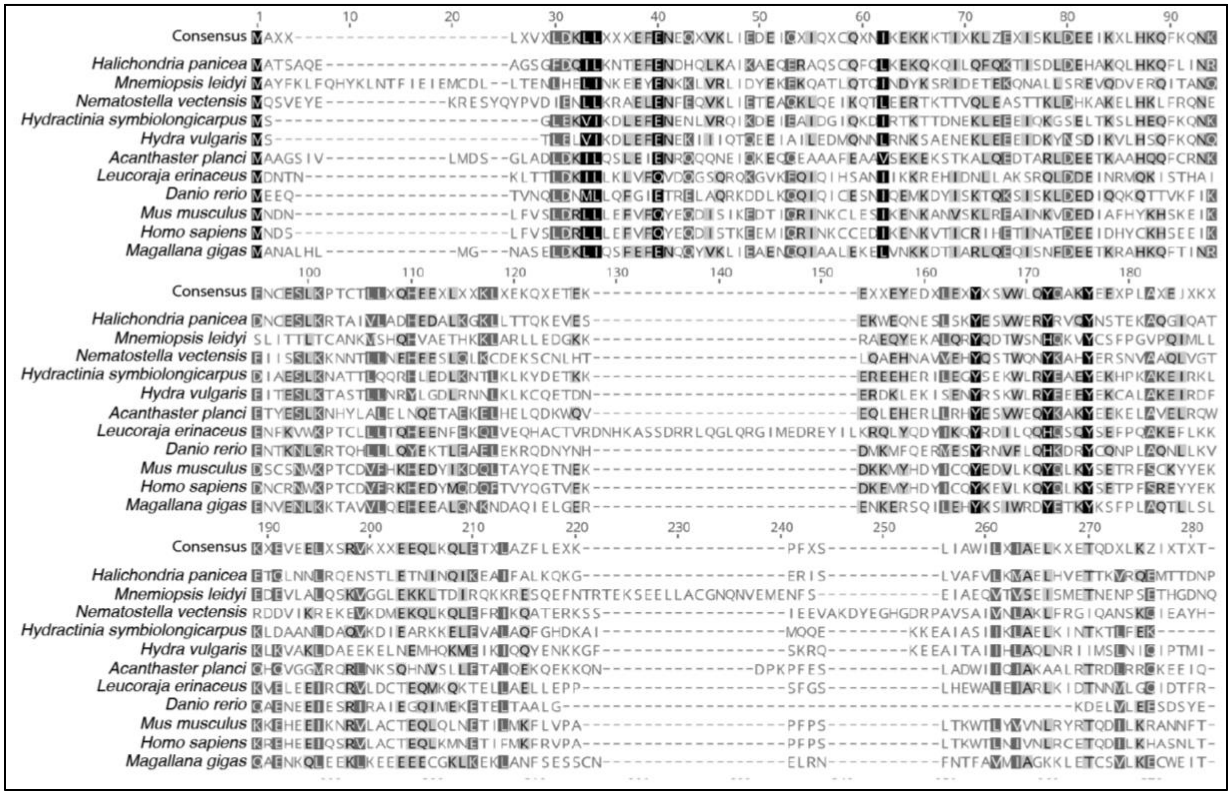

**Fig. S4. Alignment of SIX6OS1 region from various metazoan species.**

Multiple sequence alignment of SIX6OS1 N-terminal sequences. Black shading of amino acids indicates 100% residue similarity. Dark grey shading of amino acids indicates 80-99% residue similarity. Light grey shading indicates 60-79% residue similarity. White shading indicates less than 60% similarity.

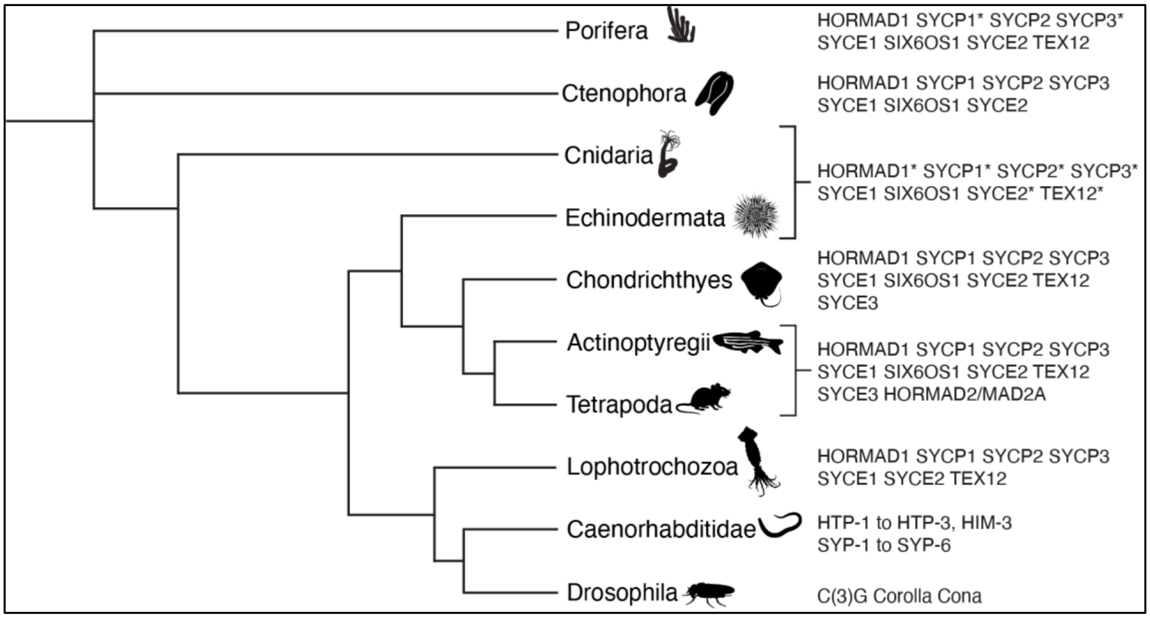


**Fig. S5. Schematic phylogenetic tree of SC proteins in different animal groups**

Schematic representation of animal groups and their phylogenetic relationships to each other. Illustrations of representative animals of each group were taken from PhyloPic, except the *Nematostella* illustration, which is from the authors themselves. SC proteins found in each group, or several groups as marked by brackets, are listed on the right. Orthologs in basal metazoan groups marked with an asterisk were previously already identified in the respective group.

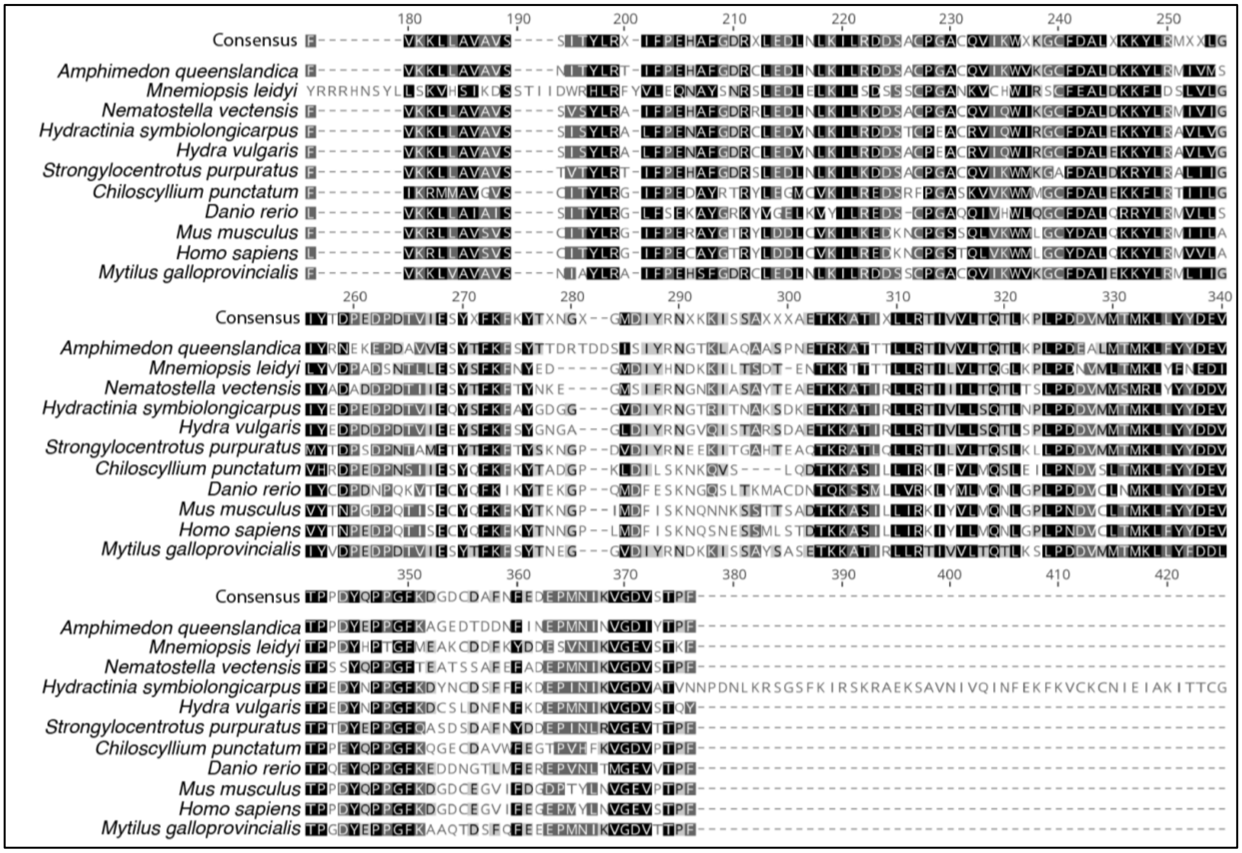


**Fig. S6. Alignment of HORMAD1 proteins from various metazoan species.**

Multiple sequence alignment of HORMA domain of HORMAD1 protein sequences. Black shading of amino acids indicates 100% residue similarity. Dark grey shading of amino acids indicates 80-99% residue similarity. Light grey shading indicates 60-79% residue similarity. White shading indicates less than 60% similarity.

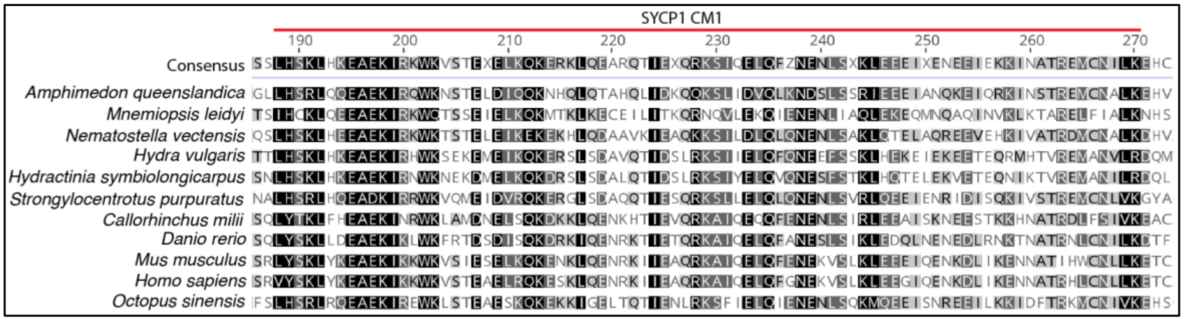


**Fig. S7. Alignment of SYCP1 conserved region 1 from various metazoan species.**

Multiple sequence alignment of SYCP1 CM1 sequences. Black shading of amino acids indicates 100% residue similarity. Dark grey shading of amino acids indicates 80-99% residue similarity. Light grey shading indicates 60-79% residue similarity. White shading indicates less than 60% similarity.

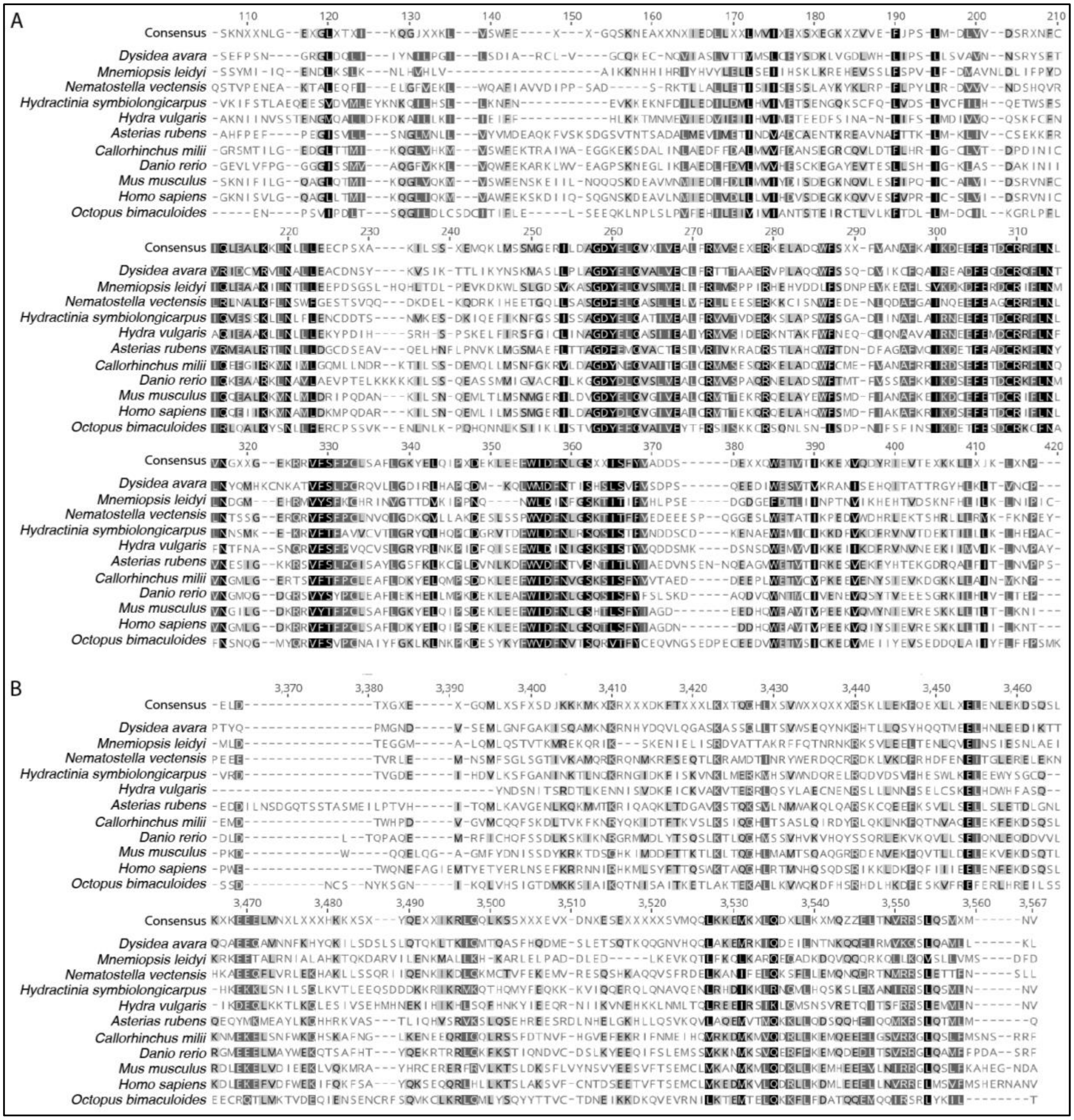

**Fig. S8. Alignment of conserved SYCP2 regions from various metazoan species.**

Multiple sequence alignment of **(A)** N-terminal and **(B)** C-terminal SYCP2 sequences. Black shading of amino acids indicates 100% residue similarity. Dark grey shading of amino acids indicates 80-99% residue similarity. Light grey shading indicates 60-79% residue similarity. White shading indicates less than 60% similarity.

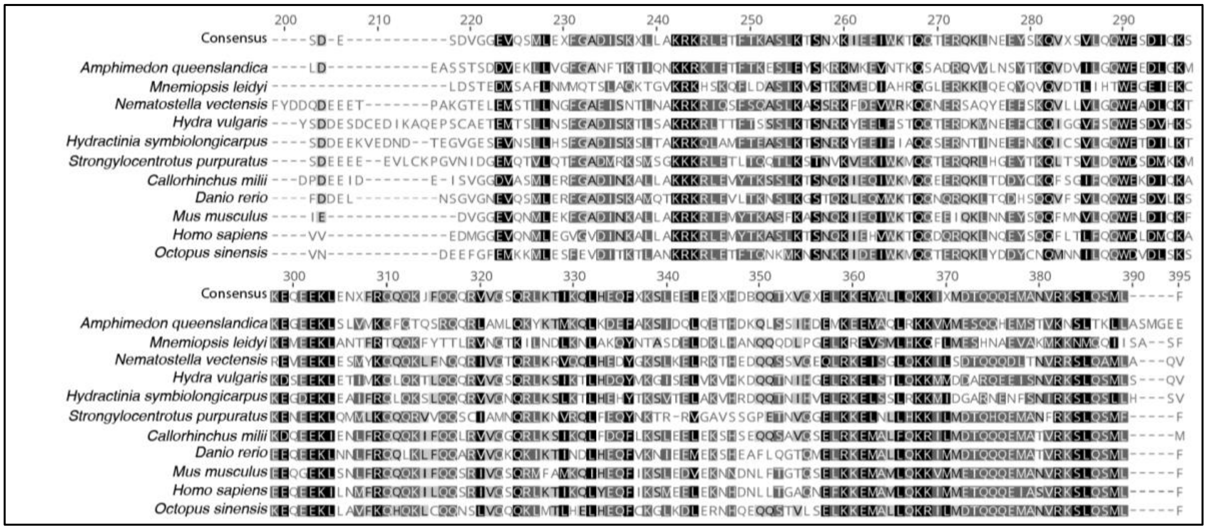


**Fig. S9. Alignment of conserved SYCP3 region from various metazoan species.**

Multiple sequence alignment of the C-terminus of SYCP1 sequences. Black shading of amino acids indicates 100% residue similarity. Dark grey shading of amino acids indicates 80-99% residue similarity. Light grey shading indicates 60-79% residue similarity. White shading indicates less than 60% similarity.

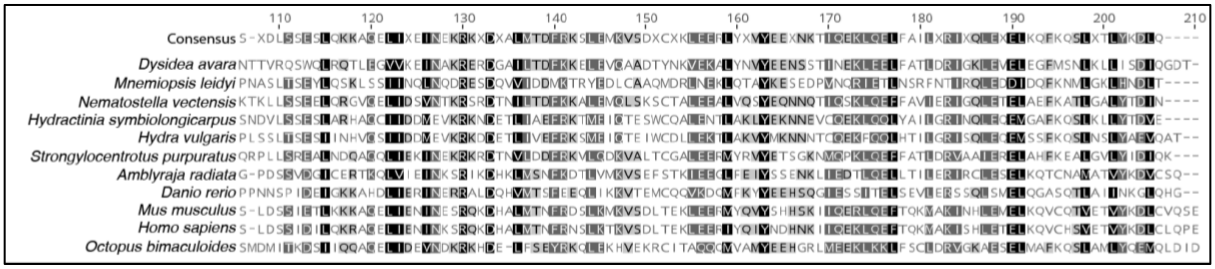

**Fig. S10. Alignment of SYCE2 region from various metazoan species.**

Multiple sequence alignment of central region of SYCE2 sequences. Black shading of amino acids indicates 100% residue similarity. Dark grey shading of amino acids indicates 80-99% residue similarity. Light grey shading indicates 60-79% residue similarity. White shading indicates less than 60% similarity.

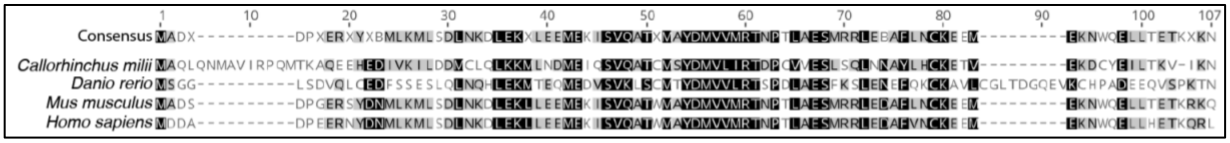


**Fig. S11. Alignment of SYCE3 from vertebrate species.**

Multiple sequence alignment of full SYCE3 sequences. Black shading of amino acids indicates 100% residue similarity. Dark grey shading of amino acids indicates 80-99% residue similarity. Light grey shading indicates 60-79% residue similarity. White shading indicates less than 60% similarity.

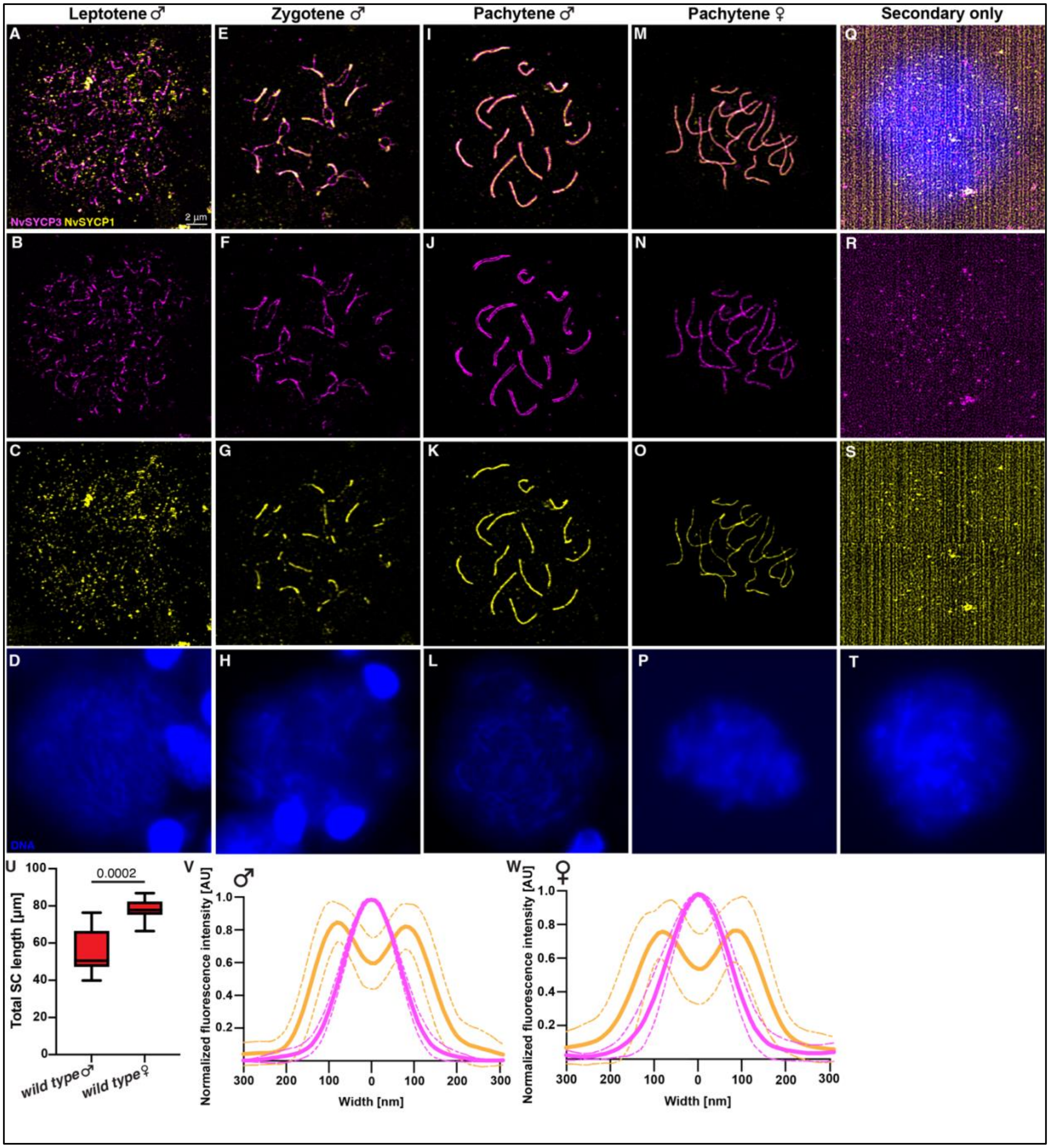


**Fig. S12. Merged and single channel images from Fig. 2**

**(A) - (T)** Merged and single channel images. All images are to the same scale. Scale bar in (A) represents 2 µm. (Q) - (T) are secondary antibody only controls. **(U)** Quantification of the total SC lengths of male and female wild type *Nematostella*. P value is from a Mann Whitney test comparing the median of these two. **(V)-(W)** Normalized fluorescence intensities of anti-SYCP1 and anti-SYP3 signal line profiles from *wild-type* pachytene SCs of males (U) and females (V).

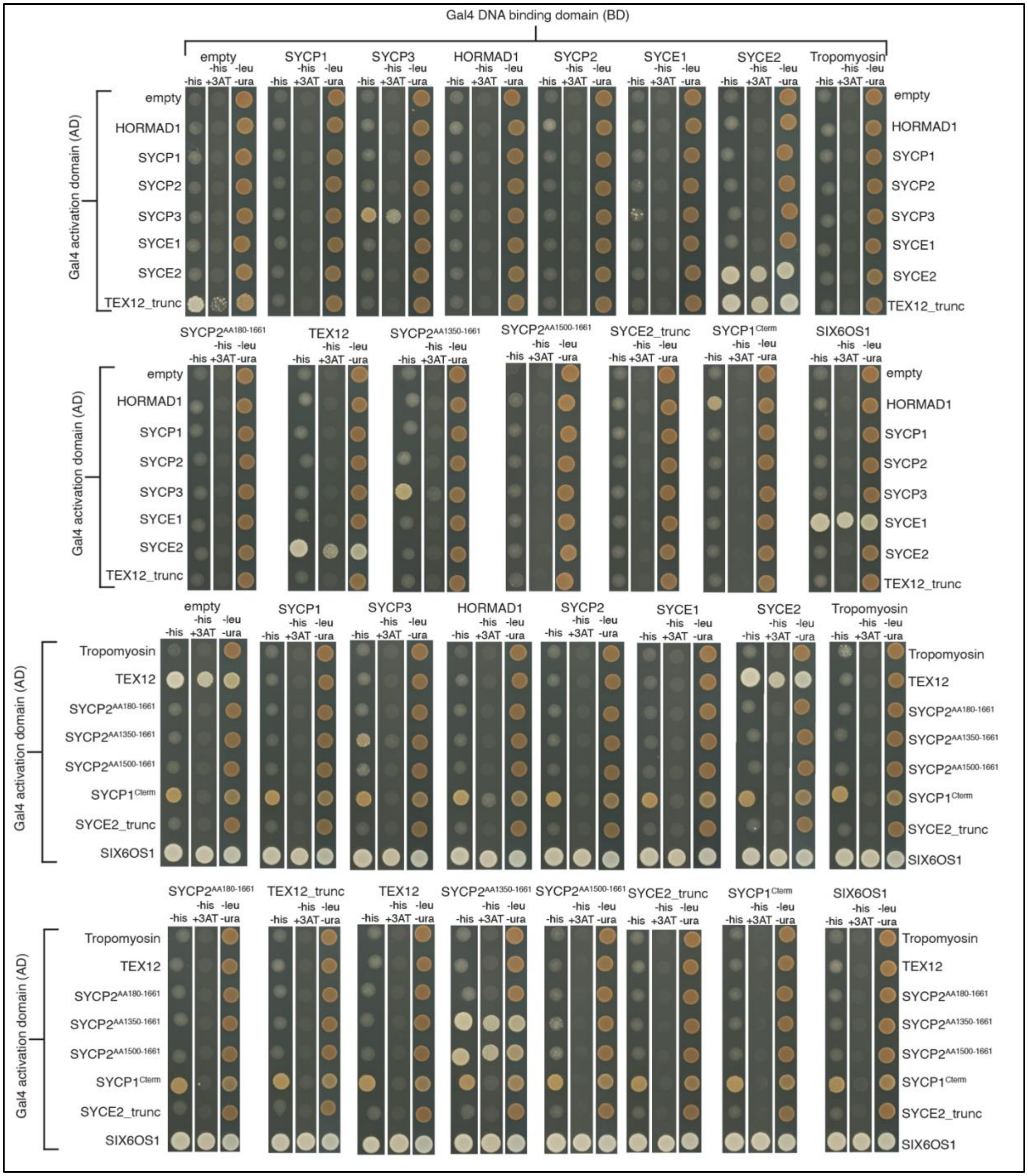


**Fig. S13. Yeast two-hybrid results of *Nematostella* SC proteins.**

Assays testing all currently known *Nematostella* SC proteins with each other for potential protein-protein interactions. Empty vector tests were done to test for auto-activation. Tests with Tropomyosin function to test whether any proteins just interact with any coiled-coil protein sequence.

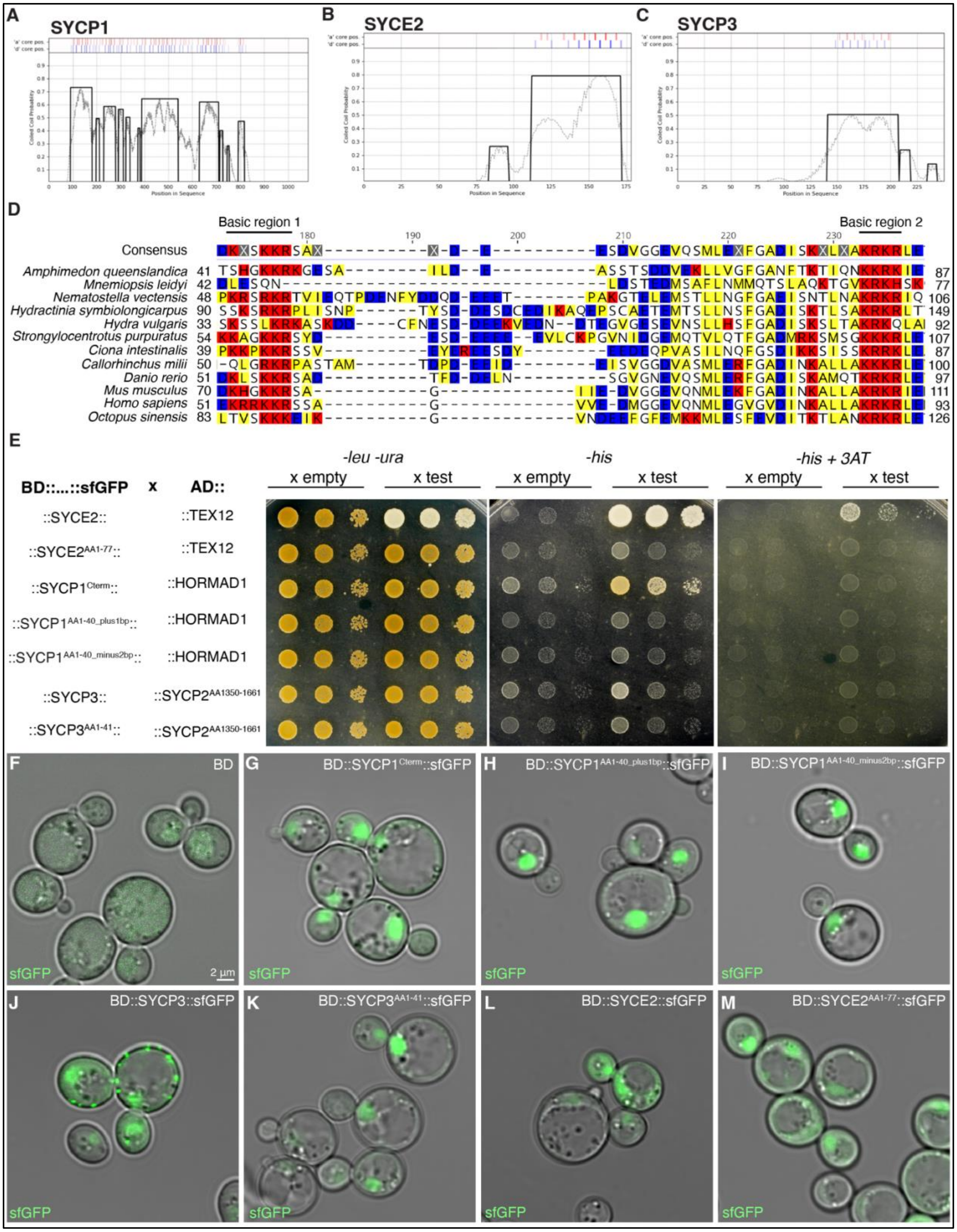


**Fig. S14. Additional information to the mutated SC proteins**

**(A) - (C)** Coiled-coil domains of SYCP1, SYC2, and SYCP3 as determined by DeepCoil2. **(D)** Multiple sequence alignment of SYCP3 amino acid sequences from different animal groups showing the conservation of two basic amino acid regions that have been shown to interact with DNA. Shading of amino acids goes as follows: red = positive charge, blue = negative charge, yellow = lipophilic. **(E)** Yeast two-hybrid results from testing wild-type and truncated proteins of interest with their previously identified protein-protein interaction partners from Fig. S13. These are additionally tagged with sfGFP to verify expression of these constructs. **(F) - (M)** Verification of sfGFP expression in yeast cells from an empty BD vector (F) and all tested constructs in (E).

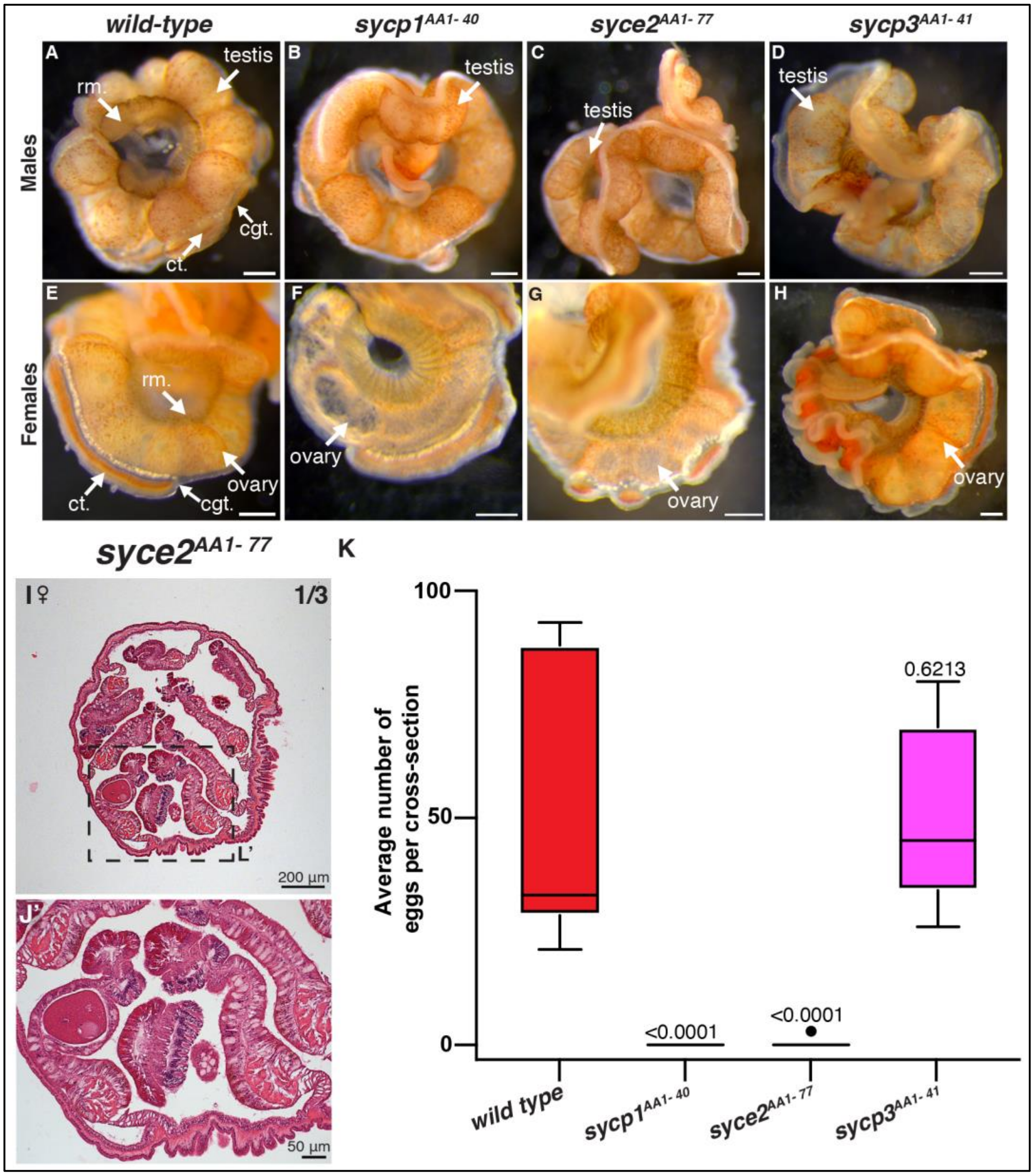


**Fig. S15. Gametogenesis in *wild-type* and mutant *Nematostella***

**(A) - (H)** Dissected mesenteries from wild-type and mutant animals. Scale bars represent 200 µm. **(I)** H&E stained cross-section of the gonadal region of a *syce2$^{AA1-77}$* female showing one egg inside one gonad. Number (1/3) indicates that only one out of three similarly examined females showed presence of eggs. Dashed line box indicates where magnified image was taken form. Scale bar represents 200 µm. **(J')** Magnified image from (I). Scale bar represents 50 µm. **(K)** Quantification of eggs found in 9 cross-sections from female gonadal regions. P values are from a Mann Whitney test comparing the median of the respective mutant to the wild-type number.

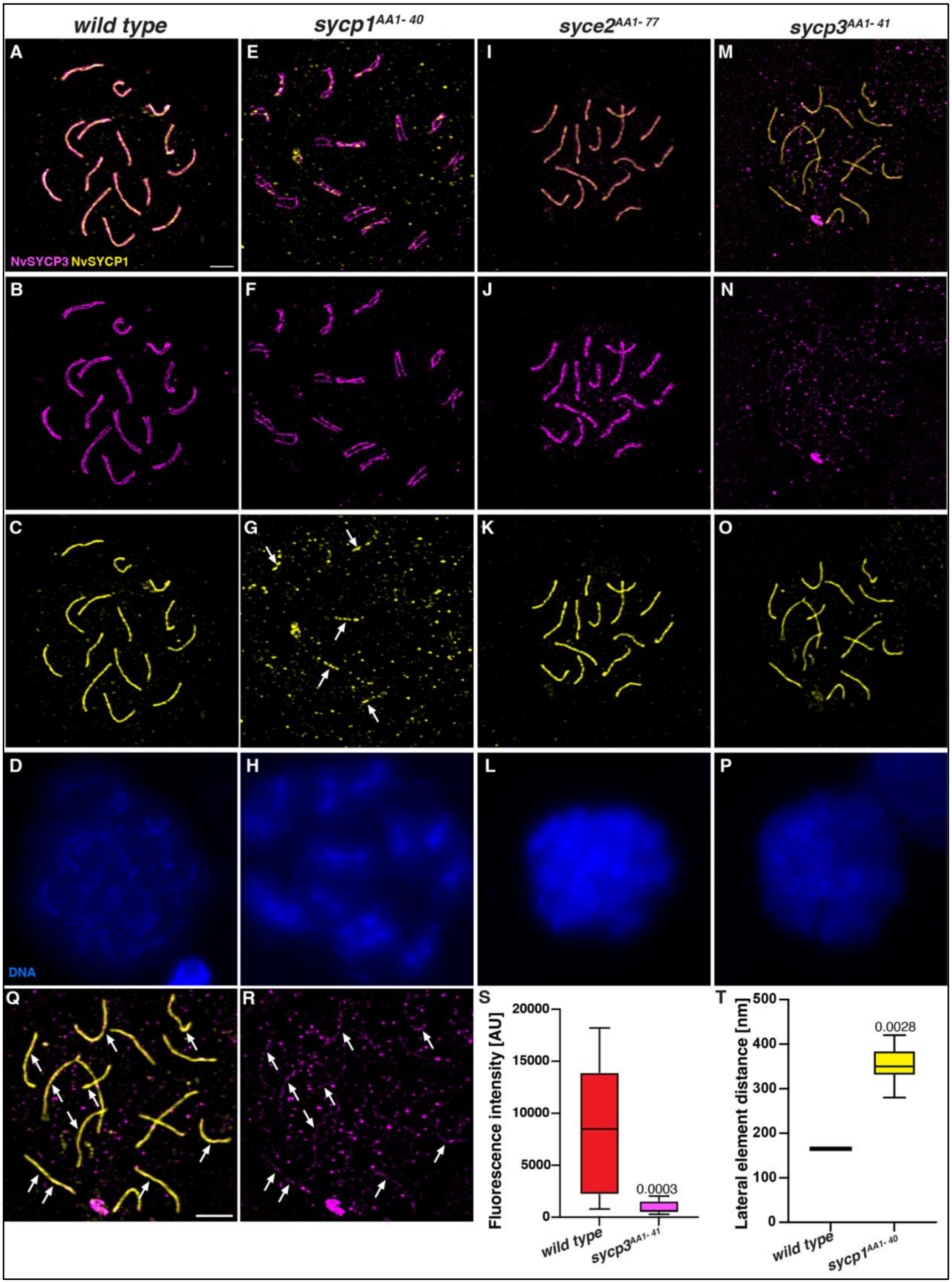
wild type
sycp1AA1-40
syce2AA1-77
sycp3AA1-41
A
E
I
M
NvSYCP3 NvSYCP1
B
F
J
N
C
G
K
O
D
H
L
P
DNA
Q
R
S
T
Fluorescence intensity [AU]
20000
15000
10000
5000
0
0.0003
wild type
sycp3AA1-41
Lateral element distance [nm]
500
400
300
200
100
0
0.0028
wild type
sycp1AA1-40

**Fig. S16. Merged and single channel images from males in Fig. 4.**

**(A) - (P)** All images are to the same scale. (G) Arrows point to SYCP1 signal indicating localization of the truncated SYCP1 N-terminus to the SC. Scale bar in (A) represents 2 μm. **(Q) - (R)** Arrows point to locations in the *sycp3*$^{AA1\text{-}41}$ male spread where low SYCP3 signal can be found. Scale bar represents 2 μm. **(S)** Quantification of the relative fluorescence intensity found in the SYCP3 channel in controls and *sycp3*$^{AA1\text{-}41}$ males. P value is from a Mann Whitney test comparing the median of the respective mutant to the *wild-type* number. **(T)** Distance between lateral elements in *wild-type* and *sycp1*$^{AA1\text{-}40}$ males. P values is from a Mann Whitney test comparing the median of the distance in *wild-type* and the *sycp1*$^{AA1\text{-}40}$ males.

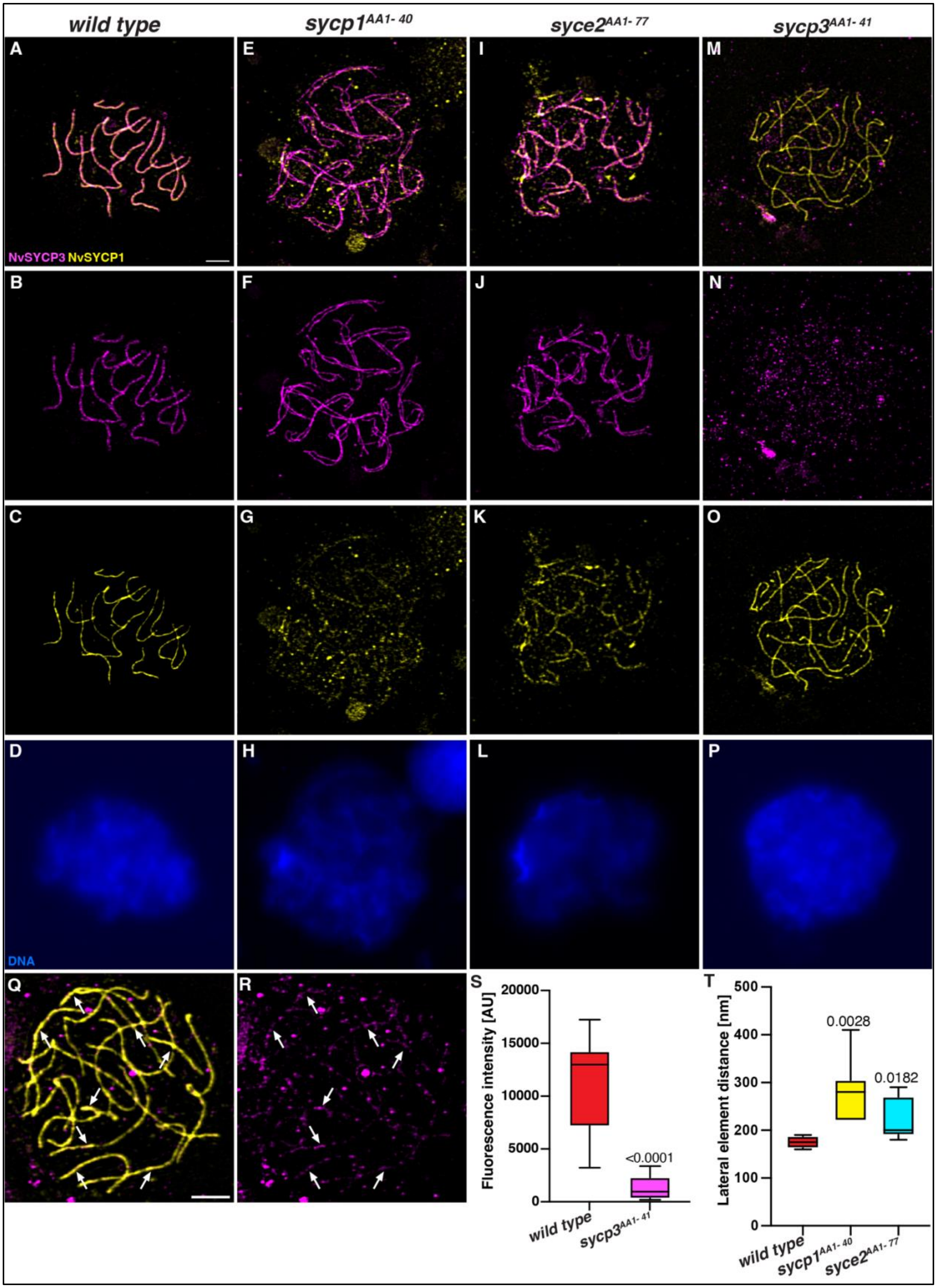
wild type
sycp1AA1-40
syce2AA1-77
sycp3AA1-41
A
B
C
D
E
F
G
H
I
J
K
L
M
N
O
P
Q
R
S
T
NvSYCP3 NvSYCP1
DNA
Fluorescence intensity [AU]
20000
15000
10000
5000
0
<0.0001
wild type
sycp3AA1-41
Lateral element distance [nm]
500
400
300
200
100
0
0.0028
0.0182
wild type
sycp1AA1-40
syce2AA1-77

**Fig. S17. Merged and single channel images from females in Fig. 4.**

**(A) - (P)** All images are to the same scale. Scale bar in (A) represents 2 μm. **(Q) - (R)** Arrowheads point to locations in the *sycp3*$^{AA1\text{-}41}$ females where low SYCP3 signal can be found. Scale bar represents 2 μm. **(S)** Quantification of the relative fluorescence intensity found in the SYCP3 channel in controls and *sycp3*$^{AA1\text{-}41}$ females. P value is from a Mann Whitney test comparing the median of the respective mutant to the wild-type number. **(T)** Distance between lateral elements in wild type, *sycp1*$^{AA1\text{-}40}$ and *syce2*$^{AA1\text{-}77}$ females. P values is from a Mann Whitney test comparing the median of the distance in the mutant females compared to the wild type.

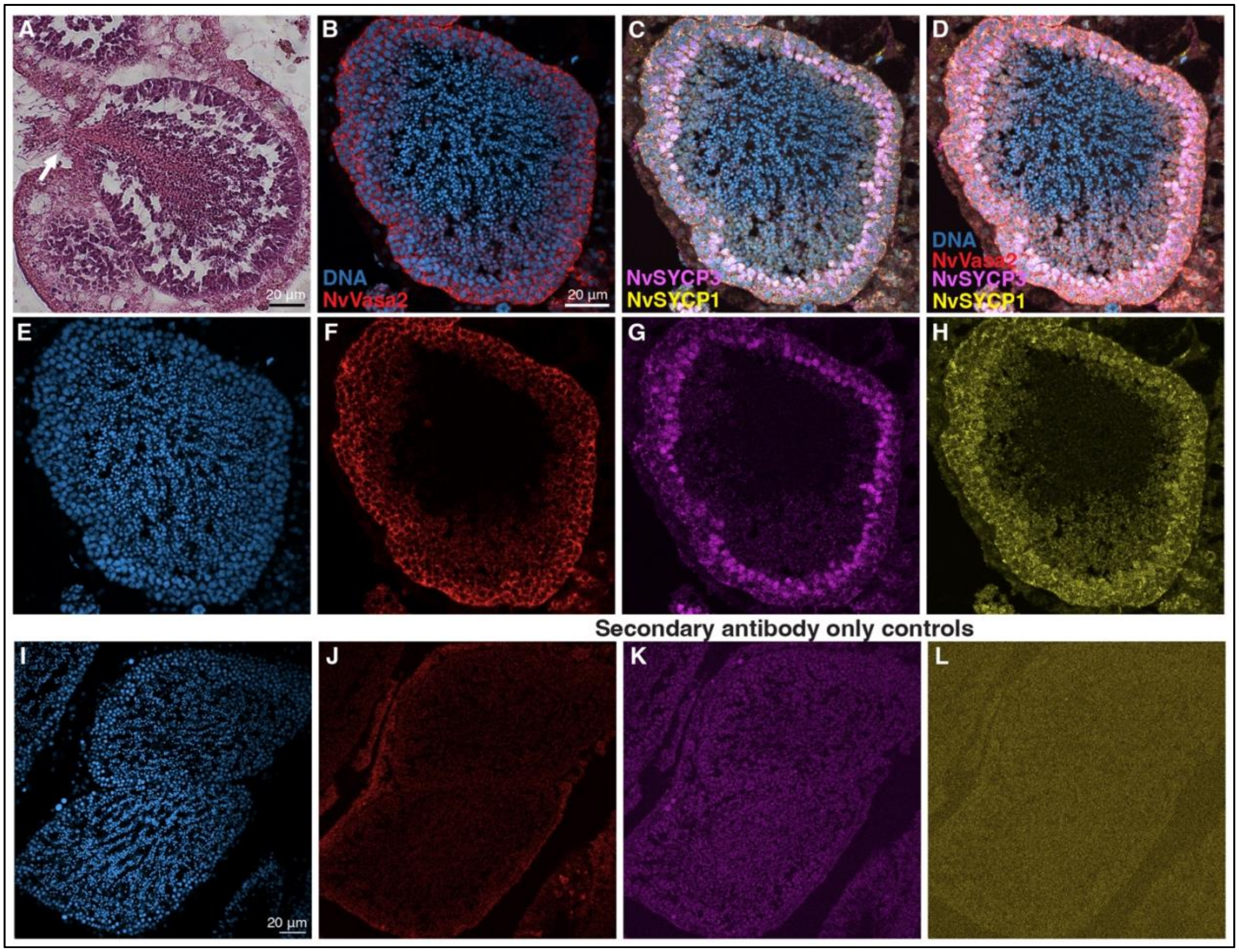


**Fig. S18. Organization of *Nematostella* spermaries.**

**(A)** Close up of a spermary in a cross-section of a male gonadal region, stained with H&E. The white arrow marks a site of sperm released from the gonad into the body lumen, visible as sperm breaking through a hole in the gonadal epithelium. Scale bar represents 20 μm and is only applicable to this individual image. **(B)-(D)** Immunofluorescence images of a spermary from a male gonad. All images are to the same scale, scale bas in (B) represents 20 μm. (B) Merged image of DNA and NvVasa2 staining. (C) Merged image of DNA, NvSYCP3, and NvSYCP1 staining. (D) Merged image of DNA, NvVase2, NvSYCP3, NvSYCP1 staining. **(E)-(H)** Individual channel images. **(I)-(L)** Hoechst with secondary-antibody-only controls on two *Nematostella* spermaries.

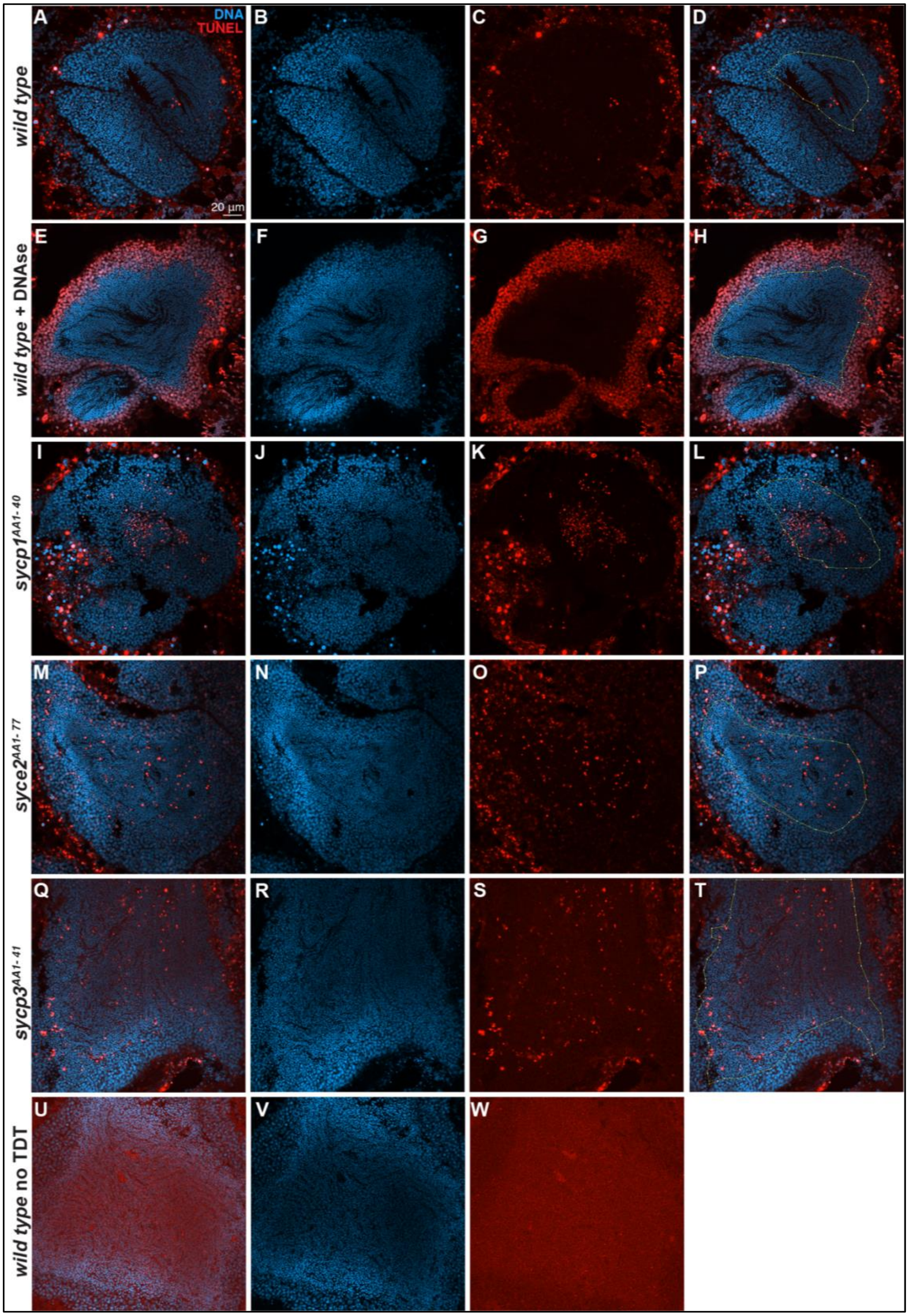
DNA
TUNEL
A
B
C
D
20 µm
E
F
G
H
I
J
K
L
M
N
O
P
Q
R
S
T
U
V
W
wild type
wild type + DNAse
sycp1^AA1-40
syce2^AA1-77
sycp3^AA1-41
wild type no TDT

**Fig. S19. TUNEL stained cross-sections of males.**

**(A) - (W)** Merged and single-channel images of Hoechst and TUNEL staining of cross-sections from *wild-type* and mutant males. Controls are wild-types exposed to DNAse I and a no TdT control. All images are to the same scale. Images **(D), (H), (L), (P), (T)** show examples of the region of interest (ROI) used for the quantification of TUNEL and DNA staining colocalization.

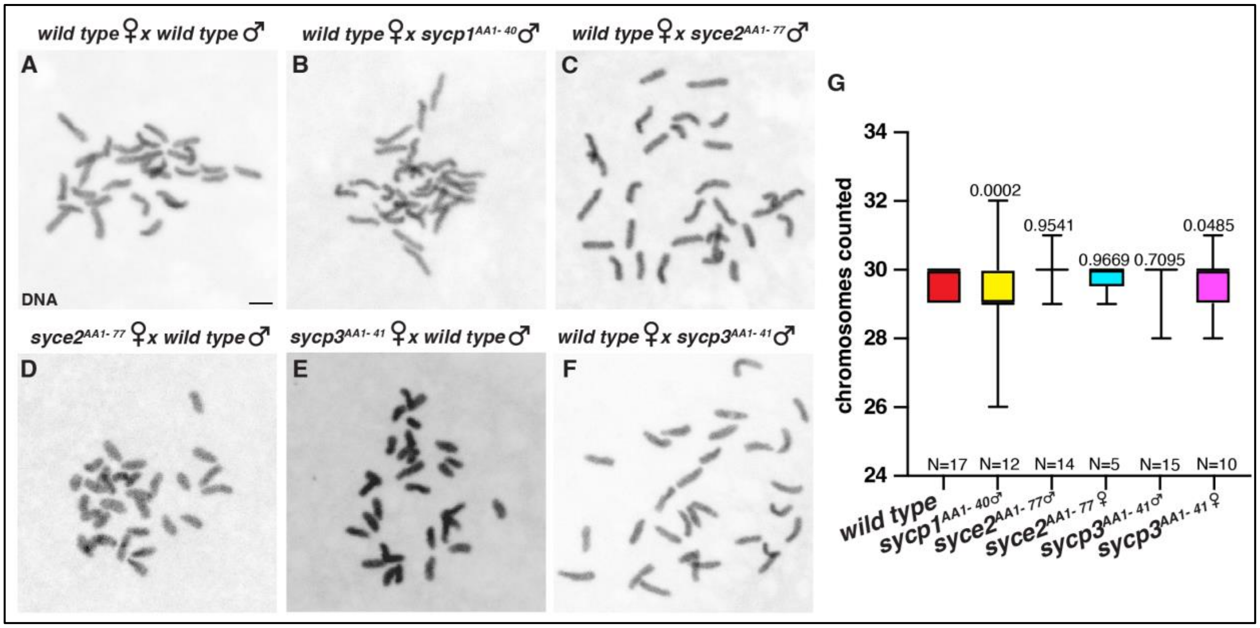


**Fig. S20. Mitotic chromosome squashes from embryos.**

**(A) - (F)** Representative mitotic chromosome squashes from embryos that resulted from the fertilization of mutant sperm/eggs with *wild-type* eggs/sperm respectively. Scale bars represent 2 μm. **(G)** Quantification of chromosomes counted in the specified number (N) of chromosome squashes. P values on top of box plots are from F tests to compare variances between the specified mutant and the control.

| Measured distance between lateral elements |
|---|
| 95 nm |
| 113 nm |
| 87 nm |

**Table S1.**

Measured distances between lateral elements of three SCs as visualized through electron microscopy.

| | *sycp3*$^{AA1-41}$ | | *sycp1*$^{AA1-40}$ | | *syce2*$^{AA1-77}$ | |
|---|---|---|---|---|---|---|
| | female | male | female | male | female | male |
| wild type | 4/4 (100%) | 23/23 (100%) | 5/5 (100%) | 5/5 (100%) | 13/13 (100%) | 13/13 (100%) |
| Heterozygote | 4/4 (100%) | 14/14 (100%) | 8/8 (100%) | 15/15 (100%) | 22/22 (100%) | 42/42 (100%) |
| homozygote | 5/5 (100%) | 7/7 (100%) | 0/6 (0%) | 7/7 (100%) | 1/20 (0.05%) | 10/10/ (100%) |

**Table S2. Spawning data from the F2 generation.**
Data listed is after all animals reached one year of age.

| | *sycp1*$^{AA1-40}$ | | *syce2*$^{AA1-77}$ |
|---|---|---|---|
| | female | male | female |
| Individual 1 | 1 | 5 | 0 |
| Individual 2 | 3 | 4 | 0 |
| Individual 3 | 3 | 4 | 3 |
| Individual 4 | N.A. | N.A. | 1 |

**Table S3. Total spreads with visible crossover sites**

Number of total spreads with visible crossover sites, as direct connections between well-separated lateral elements stained by SYCP3.
